# Dark aerobic sulfide oxidation by anoxygenic phototrophs in anoxic waters

**DOI:** 10.1101/487272

**Authors:** Jasmine S. Berg, Petra Pjevac, Tobias Sommer, Caroline R.T. Buckner, Miriam Philippi, Philipp F. Hach, Manuel Liebeke, Moritz Holtappels, Francesco Danza, Mauro Tonolla, Anupam Sengupta, Carsten J. Schubert, Jana Milucka, Marcel M.M. Kuypers

## Abstract

**SUMMARY:** Anoxygenic phototrophic sulfide oxidation by green and purple sulfur bacteria (PSB) plays a key role in sulfide removal from anoxic shallow sediments and stratified waters. Although some PSB can also oxidize sulfide with nitrate and oxygen, little is known about the prevalence of this chemolithotrophic lifestyle in the environment. In this study, we investigated the role of these phototrophs in light-independent sulfide removal in the chemocline of Lake Cadagno. Our temporally resolved, high-resolution chemical profiles indicated that dark sulfide oxidation was coupled to high oxygen consumption rates of ~9 μM O_2_·h^−1^. Single-cell analyses of lake water incubated with ^13^CO_2_ in the dark revealed that *Chr. okenii* was to a large extent responsible for aerobic sulfide oxidation and it accounted for up to 40 % of total dark carbon fixation. The genome of *Chr. okenii* reconstructed from the Lake Cadagno metagenome confirms its capacity for microaerophilic growth and provides further insights into its metabolic capabilities. Moreover, our genomic and single-cell data indicated that other PSB grow microaerobically in these apparently anoxic waters. Altogether, our observations suggest that aerobic respiration may not only play an underappreciated role in anoxic environments, but also that organisms typically considered strict anaerobes may be involved.

**ORIGINALITY-SIGNIFICANCE STATEMENT:** This study reveals that dark aerobic sulfide oxidation within an anoxic layer dominated by anoxygenic phototrophic bacteria in the stratified water column of Lake Cadagno is to a large extent carried out by the anoxygenic phototrophic bacterium *Chromatium okenii*. Our findings imply that aerobic metabolisms may be more prevalent in anoxic zones than previously thought. We also present an environmental metagenome-assembled genome of *Chr. okenii* which is the first genome sequence for the genus *Chromatium* and reveals new interesting physiological features of this environmentally relevant organism, including its capacity for aerobic respiration.

## INTRODUCTION

Anoxygenic phototrophic bacteria oxidizing sulfide and fixing CO_2_ with sunlight play an important role in the carbon and sulfur cycles of sulfidic, shallow sediments and stratified water columns. Phototrophic sulfur bacteria, for example, are believed to be responsible for 20-85% of the total daily carbon fixation in anoxic lakes (summarized in Cohen *et al*., 1977). This primary production is so important that it can control the bulk C-isotope fractionation in the water column, generating isotopic signatures that are transported and preserved in sediments (Posth *et al*., 2017). Biomass from anoxygenic phototrophs both feeds grazing zooplankton in overlying oxic waters (Sorokin 1966) and drives sulfate reduction in anoxic waters below (Pfennig 1975). The phototrophic sulfur bacteria also remove toxic sulfide from the water column, enabling aerobic life at the surface while recycling sulfur compounds for sulfate reducers. While their role in sulfide detoxification has primarily been studied in stratified lakes, there are a few examples of marine environments such as the Black Sea (Jørgensen *et al*., 1991; Overmann *et al*., 1992; Manske *et al*., 2005; Marschall *et al*., 2010) and the Chesapeake Bay (Findlay *et al*., 2015) where phototrophic sulfur bacteria also significantly impact sulfur cycling.

Anoxygenic phototrophs generally inhabit illuminated, anoxic, reducing environments which has been attributed to the toxicity of oxygen to these bacteria, and to the competition with abiotic reactions involving oxygen for their electron donors. Abiotic sulfide oxidation with oxygen is kinetically hampered and thus extremely slow (on the order of days) in trace-metal poor waters (e.g. Millero 1986; Millero et al 1987; Luther et al 2011). However, the presence of metals such as Fe, and less importantly Mn, has been shown to catalyze the reaction between sulfide and oxygen (Vazquez et al 1989). Nonetheless, sulfide oxidation by microorganisms possessing enzymes which have evolved to overcome these kinetic constraints likely remains the most important sulfide removal process in most environments (Luther et al. 2011).

In fact, some anoxygenic phototrophs have evolved the capacity for chemotrophic growth under microoxic conditions. Whereas the green sulfur bacteria (GSB) of the *Chlorobiaceae* family are considered strict anaerobes, members of the *Proteobacteria* collectively known as the purple sulfur bacteria (PSB) can be facultatively microaerobic (e.g. Kampf and Pfennig, 1980; de Witt and Van Gemerden, 1990). Both the GSB and PSB are well adapted to fluctuating environmental conditions, synthesizing and accumulating storage compounds during periods of substrate/nutrient excess. The anoxygenic phototrophs are known to store zero-valent sulfur (S^0^), polyphosphate, glycogen, and in the case of the PSB alone, poly-3-hydroxyalkanoates (PHA) (Mas and Van Gemerden, 1995). The macromolecular structure and metabolism of these compounds have been intensely studied in laboratory pure cultures in order to understand conditions leading to their accumulation and breakdown. It has been suggested that glycogen may play a role in energy generation under dark conditions based on observations that cultured *Chromatium* sp. utilize glycogen to reduce stored sulfur, yielding sulfide and PHA (Van Gemerden, 1968).

Here we investigated the role of anoxygenic phototrophic bacteria in dark sulfur cycling processes in Lake Cadagno, a permanently stratified lake with high sulfate concentrations of up to 1-2 mM in the monimolimnion. Microbial reduction of sulfate produces large amounts of sulfide in the anoxic bottom waters (up to ~300 μM) and sediments (>500 μM) which support dense populations of GSB and PSB in the photic zone. These bacteria heavily influence the chemistry of the lake, forming chemocline of 1-2 meters in thickness where sulfide and oxygen remain mostly non-detectable. The PSB *Chromatium okenii* is by far the most active of these bacteria, having been shown to play a disproportionately large role in inorganic carbon and ammonium assimilation despite their low abundances (<1% of total cell numbers) in the chemocline (Musat *et al*., 2008; Posth *et al*., 2017). In addition to their important contribution to light-driven sulfide oxidation, previous studies have shown that the anoxygenic phototrophic bacteria of Lake Cadagno remain active in the dark (Musat *et al*., 2008; Halm *et al*., 2009; Storelli *et al*., 2013). However, their mechanism of energy generation in the absence of light is not yet clear. We therefore combined high-resolution biogeochemical profiling with metagenomic analyses to gain an overview of possible light-independent metabolic processes impacting the sulfur biogeochemistry of Lake Cadagno. In addition to providing insights into the metabolism of anoxygenic phototrophic bacteria *in situ*, we present a model to explain the mechanism of dark sulfide oxidation in the chemocline of this meromictic lake.

## RESULTS & DISCUSSION

### Biogeochemistry of Lake Cadagno

Lake Cadagno is characterized by an oxic mixolimnion and a sulfidic monimolimnion spatially separated from each other by a chemocline (defined by bold contour lines in Fig. 1A) free of detectable oxygen (detection limit 50-100 nM) and containing very little sulfide (0-5 μM). In August 2015, oxygen disappeared just above the chemocline close to 12 m depth. The daytime increase in oxygen concentrations between 11-12 m depth denotes net photosynthesis and the nighttime decrease denotes net respiration (Fig. 1A). The permanent absence of oxygen in the chemocline indicated that oxygen was consumed both in the day and the night.

**Fig 1.**
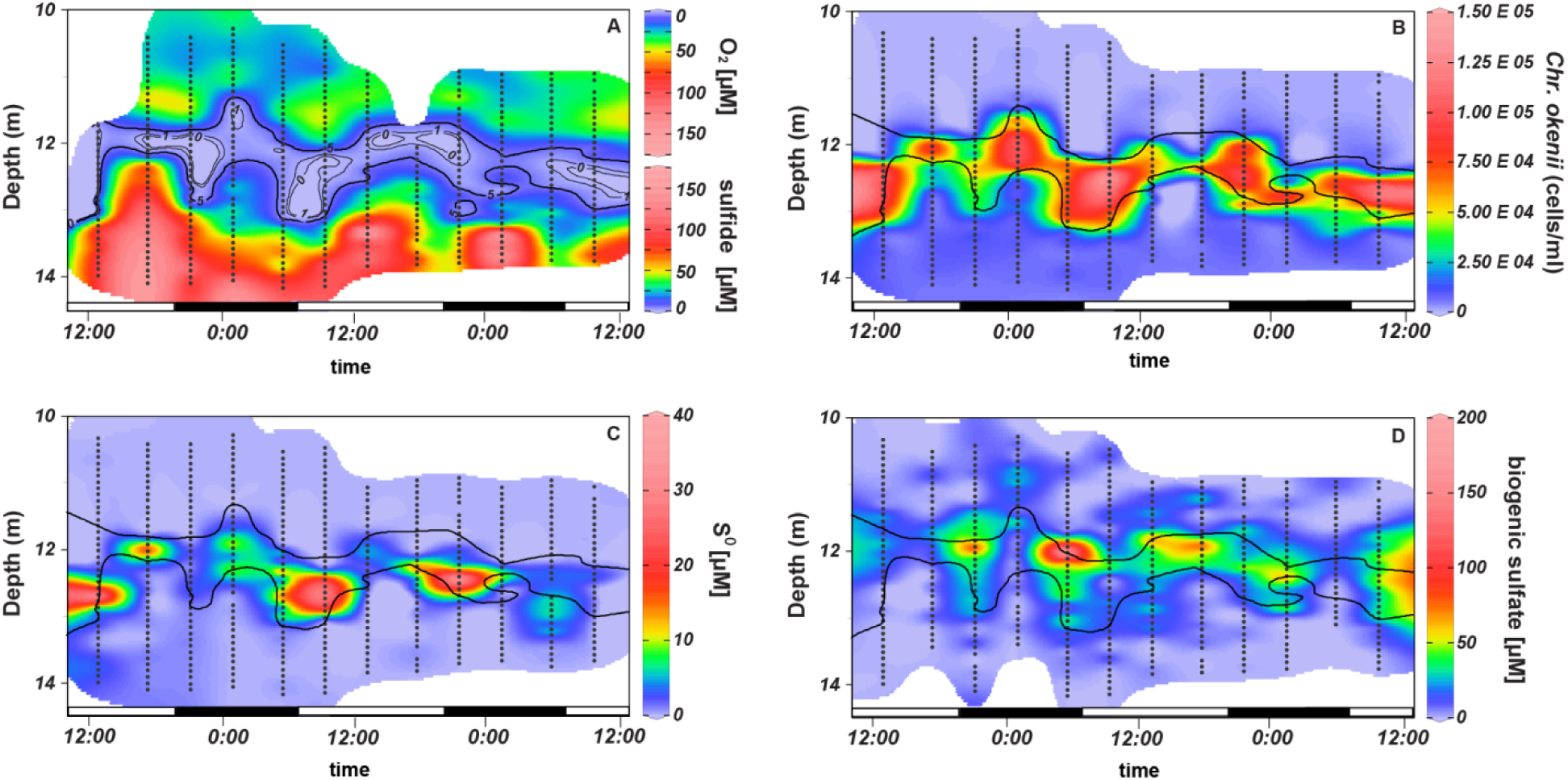
(A) Combined oxygen (top) and sulfide (bottom) profiles of the Lake Cadagno water column revealing the persistence of an oxygen- and sulfide- free zone over a period of 48 hours, with contour lines indicating sulfide concentrations. The bold contour lines delimiting the region with > 5 μM sulfide were used to define the chemocline in parallel profiles of *Chr. okenii* cell counts (B), particulate S^0^ (C), and sulfate (D). Black dots represent sampling points for all parameters except O_2_ which was measured with a microsensor mounted on a CTD probe. Shaded boxes represent dark periods between sunset at ~20:50 and sunrise at ~6:10. Time plots were interpolated from original profiles measured in August 2015 and are provided in Fig S1.

Steep gradients of sulfide diffusing into the chemocline varied independently of light-dark periods and the total sulfide concentration in the chemocline did not exceed 5 μM at any time point. Because the lake is meromictic, these stratified conditions were also present during other sampling years (see Fig. S2 for 2013 and 2014 profiles). In 2015, the 0.5-1 m wide chemocline was located around 11-12 m depth, with the exact location varying over the day most likely due to the action of internal waves (Egli *et al*., 1998). In previous years, the chemocline was up to 2 m wide (Fig. S2) and remained completely sulfide-free in the dark. Conservative properties such as temperature and conductivity were constant throughout the chemocline in all years sampled (Fig. S1&2) indicating mixing of this zone (Sommer *et al*., 2017). Flat conductivity profiles revealed stronger mixing of the chemocline in 2013 and 2014 (Fig. S2) than in 2015 (Fig. S1) when the region of constant conductivity was reduced or absent.

*Chr. okenii* was the most significant microorganism in the chemocline both in terms of biomass, accounting for ~60-80% of total microbial biovolume (Sommer *et al*. 2017), and activity, accounting for 30-40% of inorganic carbon fixation (Fig. S3). The cell abundances of *Chr. okenii* in the Lake Cadagno chemocline were enumerated by flow cytometry during 2 daily cycles (Fig. 1B). Higher densities of *Chr. okenii* were found in 2014 (10^6^·ml^−1^) than in 2015 (10^5^·ml^−1^). *Chr. okenii* is highly motile, swimming at speeds of ~27 μm·s^−1^ and has been hypothesized to drive the convection and mixing of the chemocline (Wüest, 1994; Sommer *et al*., 2017). *Chromatium* are known to migrate between gradients of sulfide, light, and oxygen by photo- and chemotaxis (Pfennig *et al*., 1968). We observed that *Chr. okenii* were positioned between oxygen and sulfide gradients, regardless of changes in depth or light availability (Fig. 1A,B). The hourly variations in cell numbers were likely the result of their heterogeneous distribution across the chemocline as *Chr. okenii* can swim in a horizontal direction or be displaced by compression of the water masses by internal waves (Egli et al., 1998). Other anoxygenic phototrophs that have been consistently detected in the chemocline include the PSB *Lamprocystis, Thiocystis* and *Thiodictyon* and several GSB of the genus *Chlorobium* (Tonolla *et al*., 1999, 2004, 2005). Together these bacteria constituted the majority of the total phototrophic cells (10^6^·ml^−1^) in 2015, but they are considerably smaller in size (and thus proportion of total biovolume) than *Chr. okenii*.

The oxidation of sulfide by these anoxygenic phototrophs proceeds via the formation of S^0^ as an obligate intermediate (Mas and Van Gemerden, 1995). This S^0^ was measured as particulate sulfur on 0.7 μm filters and may comprise S^0^ stored intracellularly by PSB and S^0^ adhering extracellularly to GSB. The highest concentrations of S^0^ (up to 45 μM; 1C) were measured in the middle of the chemocline. It is likely that this S^0^ was present in the form of both elemental S and polysulfides formed by the reaction of free sulfide with intra- and extracellular S^0^, as has previously been suggested in other euxinic lakes (Overmann, 1997). Our analytical method for total S^0^ did not distinguish between different forms of S^0^ such as cyclooctasulfur and polysulfides. However, we could confirm the presence of polysulfides inside live *Chr. okenii* cells in environmental samples using Raman spectroscopy. The Raman spectrum of a sulfur inclusion from *Chr. okenii* exhibited two weak peaks at 152 and 218 and a prominent peak at 462 cm^−1^ (Fig. S4) which is characteristic of linear polysulfide species (Janz *et al*., 1976). The Raman peak at ~2900 cm^−1^ corresponds to the CH_2_ and CH_3_ stretching vibrations (Socrates, 2004), and its co-occurrence with polysulfide peaks support the theory that the sulfur chains in these purple sulfur bacteria are terminated by organic end groups as reported previously (Prange *et al*., 1999).

Over two diurnal cycles, the S^0^ inventory (Fig. S5A), or the total amount of particulate S^0^ in the chemocline, was much lower than expected from the sulfide gradients and corresponding sulfide fluxes (discussed below), suggesting that stored S^0^ served only as a transient intermediate and was rapidly oxidized to sulfate. No day-night trends in S^0^ accumulation were apparent in the chemocline. Nevertheless, the increase in the S^0^ inventory at several time points during the night was indicative of dark sulfide oxidation.

In culture, *Chromatium* spp. are known to store carbon compounds like glycogen and polyhydroxyalkanoates (PHAs) which have been proposed to be involved in dark sulfur metabolism (Mas and van Gemerden, 1995). We therefore quantified glycogen and PHA abundance in biomass samples from one day/night profile of the chemocline (Fig. 2A&B). We could not detect any PHA, but the presence of glycogen during the day and night coincided with *Chr. okenii* cell numbers (Fig. 2A&B). This is consistent with previous reports of glycogen storage and an absence of PHA in natural populations of *Chr. okenii* (Del Don *et al*., 1994). While the highest potential cellular glycogen content (2.38 · 10^−6^ μg/cell) was found at the top of the chemocline during the day, we observed little change in the cellular glycogen content between day and night (Fig. S6). Average potential cellular glycogen decreased from 5.50 · 10^−7^ μg/cell during the day to 5.33 · 10^−7^ μg/cell during the night, which represents a 3% reduction in cellular glycogen reserves. *Chr. okenii* in Lake Cadagno were previously reported to decrease their glycogen reserves by 50 % in the dark (Del Don *et al*., 1994), but this may have been a result of undersampling as our time- and depth-resolved biogeochemical profiles revealed light-dark independent variations in *Chr. okenii* cell numbers and glycogen concentrations. While it has been demonstrated that *Chromatium* sp. in pure cultures obtain energy from the reduction of S^0^ with glycogen in the dark (Van Gemerden, 1968), we could not confirm this observation for *Chr. okenii in situ*. From our data, we conclude that storage compounds did not play a significant role in the dark respiratory metabolism of *Chr. okenii* in the Lake Cadagno chemocline.

**Figure 2.**
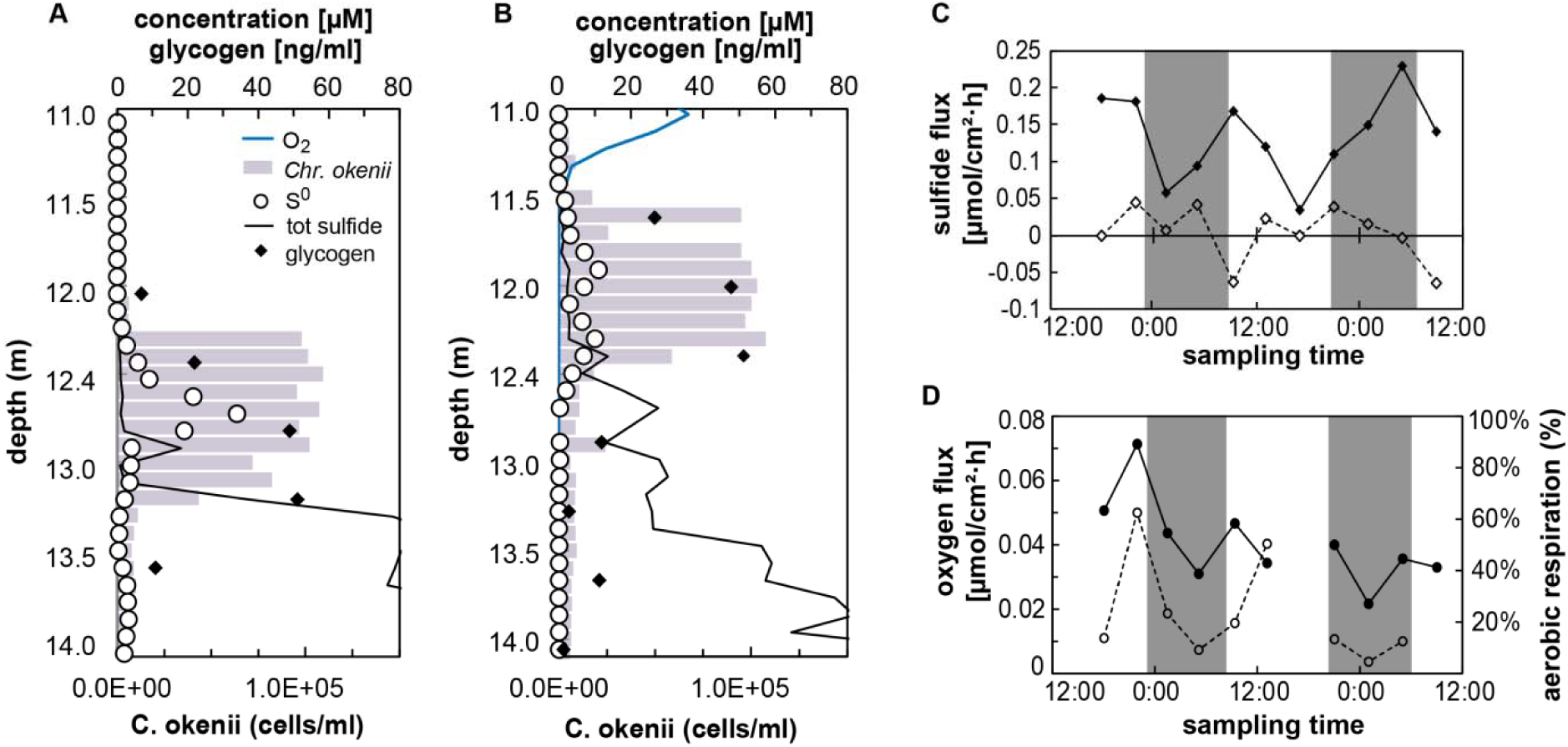
Representative day (A) and night (B) profiles through the chemocline illustrating glycogen and S^0^ concentrations in relation to *Chr. okenii* cell numbers, oxygen, and sulfide gradients in August 2015. No oxygen data is available for the day profile. Profiles were measured at 4-h intervals over 2 day-night cycles and used to calculate chemical fluxes. (C) The consumed sulfide flux (solid line) was calculated by subtracting the residual sulfide flux (dashed line) from the total sulfide flux into the mixed layer. (D) The downwards oxygen flux into the chemocline (solid line) was used to estimate the maximum % of sulfide aerobically respired (dotted line), assuming the complete oxidation of sulfide to sulfate. Shaded regions represent dark periods.

Sulfate was measured as the end product of sulfide oxidation, but due to the high (1-2 mM) background sulfate concentrations, the comparably small concentration changes resulting from sulfide oxidation processes were non-detectable. To identify regions of sulfate production in and around the chemocline, we therefore determined deviations from the sulfate-conductivity mixing line drawn for each profile (see Fig. S7 for details). Strong mixing of the chemocline is expected to produce a linear relationship between sulfate and conductivity, and large digressions from this best-fit line indicated that sulfate was produced faster than the rate of mixing. The expected sulfate concentration could be extrapolated based on measured conductivity, and then subtracted from the measured sulfate concentration to give excess sulfate:

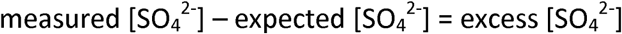

This excess sulfate was attributed to biological sulfate production, as abiotic sulfide oxidation has been shown to produce mainly S^0^ and thiosulfate both in laboratory solutions (Chen and Morris 1972; Gourmelon et al 1977) and in similarly stratified natural environments (Zopfi et al 2001; Ma et al 2006). Biogenic sulfate profiles from 2015 plotted over two diurnal cycles exhibited a peak at the top of the chemocline in the region of oxygen depletion (Fig. 1D). Interestingly, sulfate production was observed during the night and the overlap of excess sulfate and oxygen in these profiles indicated that sulfide may be oxidized aerobically. Daytime sulfate production in 2014 was related to photosynthetically active radiation (PAR) intensity (Fig. S2), suggesting that sulfide and S^0^ could either have been oxidized aerobically within the chemocline using *in situ*-produced oxygen (Milucka *et al*., 2015) or phototrophically. The comparatively broad biogenic sulfate peak in the 2014 night profile likely reflects the broader vertical distribution of the *Chr. okenii* population (Fig. S2).

The sulfate excess in the chemocline is not expected to be affected by sulfate reduction as no sulfate reduction was detected within the chemocline in 2014 or 2015. The sulfate reduction rates measured in the sulfidic zone 1 m below the chemocline were about 235 nM·d^−1^ and 375 nM·d^−1^ in 2014 and 2015, respectively.

To quantify biological sulfide consumption over time, we calculated the total sulfide flux into the chemocline (Fig. S5B). Assuming that phototrophic sulfide oxidation ceases in the dark, upwards-diffusing sulfide should accumulate in the chemocline at night. The expected sulfide accumulation was calculated based on fluxes into the layer over a 10-h night period and compared to the actual sulfide concentration observed in the layer. From an average sulfide flux *F* = 0.15 μm·cm^−2^h^−1^ (Fig. S5B), into a well-mixed layer of thickness *H* = 1 m over *t* = 10 hours, the resulting sulfide concentration *C* = *F*t/H* should be about 15 μM in the chemocline. However, the average sulfide measured in the layer did not exceed 3 μM (Fig. 1A), or five times less than expected, indicating that sulfide is consumed.

We therefore partitioned the total sulfide flux into two fractions: the flux of biologically consumed sulfide and the flux of residual sulfide in the chemocline. First, the amount of residual sulfide was calculated at each sampling time point by integrating sulfide concentrations within the mixed layer (Fig. S5C). The rate of sulfide accumulation was then calculated for each 4-h sampling interval and subtracted from the total sulfide flux to give the biologically consumed sulfide flux. The flux of sulfide consumed in the dark was in the same range as in the day (0.03 to 0.22 μmol·cm^−2^h^−1^) and the residual sulfide flux was very small in comparison (Fig. 2C). The observed variations did not correlate with day-night cycles and the changes of sulfide gradients could have been induced by internal waves, as mentioned above. Together, this indicates that sulfide oxidation continued in the dark and seemed to be related to the total sulfide flux (Fig. S5 B) rather than the presence of sunlight. For comparison, the upwards flux of sulfide in previous years was slightly lower, or 0.011-0.024 μmol·cm^−2^h^−1^ in 2013 and 0.032-0.072 μmol·cm^−2^h^−1^ in 2014.

It was not possible to calculate S^0^ fluxes in Lake Cadagno because S^0^ is actively transported by the motile purple sulfur bacteria during chemo- and phototaxis (Pfennig *et al*., 1968) independent of diffusive processes. The total (upwards and downwards) biogenic sulfate flux (Fig. S5 D) in this region was roughly equivalent to the sulfide flux and followed a similar trend.

Overall, our high-resolution profiles revealed that sulfide in Lake Cadagno was consumed during the day and night, but only light-dependent sulfide oxidation has thus far been recognized as a major sulfide-removing process in the lake. In the absence of light, it is also possible that alternative electron acceptors such as 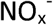, Fe^(III)^, Mn^(IV)^, or O_2_ play a role in sulfide oxidation. Nitrate and nitrite concentrations in the Lake Cadagno chemocline are negligible (Halm *et al*., 2009; Milucka *et al*., 2015). High fluxes of reduced, dissolved metals (0.027 μmol Fe·cm^−2^·d^−1^ and Mn 0.049 μmol Mn·cm^−2^·d^−1^) suggest that Fe- and Mn-oxides are rapidly reduced by microorganisms or abiotically by sulfide in the chemocline (Berg *et al*., 2016), but re-oxidation of Fe and Mn would ultimately depend on oxygen in the dark. We therefore considered oxygen as the principal direct (or indirect) oxidant responsible for observed dark sulfide oxidation.

The oxygen flux into the chemocline varied slightly between 0.022-0.071 μmol·cm^−2^h^−1^ over the period of 48 h (Fig. 2D). Oxygen fluxes measured in 2013 and 2014 were in the same range, or 0.013-0.048 μmol·cm^−2^h^−1^ and 0.037-0.073 μmol·cm^−2^h^−1^, respectively.

Sulfide oxidation can occur in two steps:

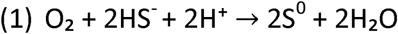

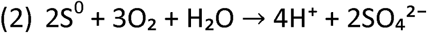

Assuming that all sulfide is eventually oxidized to sulfate, we related oxygen fluxes to sulfide consumption using the 2:1 stoichiometry for complete aerobic sulfide oxidation:

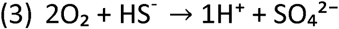

Calculated oxygen fluxes in 2013 and 2014 were sufficient to account for all the sulfide oxidized in the dark. In 2015, aerobic sulfide respiration could account for up to 10-50% of sulfide oxidized during the day and 5-45% of sulfide oxidized during the night (Fig. 2D). During the day, the remainder of sulfide oxidation could be attributed to anoxygenic photosynthesis and/or aerobic sulfide oxidation fueled by *in situ* oxygen production by photosynthetic algae. At several time points in the dark, however, we could not explain the disappearance of roughly 60-90% of upwards-diffusing sulfide. We hypothesize that the missing oxygen is supplied laterally from the turbulent transport initiated by internal wave breaking at the lake boundaries. The convection within the chemocline may be key to the transport of oxygen and sulfide to aerobic sulfide-oxidizing bacteria in the chemocline. A weakening of the mixing regime was observed in August 2015 (Sommer *et al*., 2017) which may have signified a slowed transport of electron acceptors, thus contributing to the accumulation of sulfide in the chemocline.

To determine potential O_2_ consumption rates in the Lake Cadagno chemocline, we performed incubation experiments with chemocline water by sequentially injecting dissolved O_2_ and H_2_S and monitoring O_2_ concentrations online (Fig. S8). Background rates of O_2_ consumption prior to H_2_S addition were 0.55 - 0.57 μM·h ^−1^ (Fig. S8 B). Within 5 minutes of H_2_S addition, the O_2_ consumption rate increased 4- to 5-fold and remained high for 15-30 minutes. The response of the H_2_S addition was highly reproducible, resulting in mean rates of 8.7 ± 1.9 μM O_2_·h^−1^ for five consecutive additions (Fig. S8 B). These potential aerobic respiration rates were well within the range of the volumetric oxygen consumption rates (4.4-14.2 μM O_2_·h^−1^) determined from *in situ* fluxes over a chemocline of 0.5 m thickness.

For each injection of 1.6 μM H_2_S, O_2_ concentrations fell by 1.5 to 2.5 μM which is equivalent to a stoichiometry of O_2_:H_2_S between 1:1 and 2:1. This suggests a combination of sulfide oxidation to sulfur and to sulfate (equations 1 and 3). Based on the experimental oxygen consumption rate, we calculated the most conservative sulfide oxidation rate (using a 2:1 ratio) to be 4.35 μM H_2_S·h^−1^. These rates are much higher than would be expected from abiotic sulfide oxidation alone (Millero 1986; et al. Millero 1987) implying that microorganisms play a key role in aerobic sulfide oxidation in Lake Cadagno.

### Dark activity of Chr. okenii in Lake Cadagno

While the sulfide-oxidizing *Chr. okenii* is one of the most active microorganisms in the Lake Cadagno chemocline in terms of C-fixation in the light (Musat et al. 2008; Fig. S3), its metabolic activity and contribution to sulfide removal in the dark has not yet been investigated. The capacity for microaerophilic respiration is known for a few PSB, and it is therefore possible that *Chr. okenii* grows chemoautotrophically on oxygen and sulfide in the absence of light. To test this, we incubated Lake Cadagno chemocline water with added sulfide and ^13^CO_2_ in the dark under anoxic, low microoxic, and medium microoxic conditions and compared the C-fixation activity of single *Chr. okenii* cells to that under light conditions. In order to mimic the diffusive flux of O_2_ in the lake, air was injected into the headspace of the microoxic incubations to achieve expected concentrations of 5 μM and 15 μM dissolved O_2_ in the low and medium incubations, respectively. Although bottles were shaken at the start of the incubation to equilibrate the water and headspace and then stirred constantly to enhance diffusion of O_2_ into the water, dissolved O_2_ concentrations dropped rapidly within the first hour of incubation (Fig. S9) at the same rate (8 μM O_2_·h^−1^) observed in our experimental rate measurements above. For the remainder of the incubation, dissolved O_2_ remained below 15 % of the targeted concentrations, most likely because O_2_ consumption outpaced diffusion resulting in O_2_ concentrations close to those observed *in situ*.

The incorporation of ^13^CO_2_ into *Chr. okenii* cells under dark, anoxic conditions was only slightly above background (Fig. 3A). The mean growth rate of 0.009 ± 0.007 d^−1^ could be attributed to trace oxygen contamination or to fermentation of carbon storage compounds as reported by Van Gemerden (1968). A few small rod-shaped cells were also enriched in ^13^C and were likely sulfur-disproportionating bacteria such as *Desulfocapsa* and *Desulfobulbus* which have previously been identified in Lake Cadagno (Tonolla et al. 2000; Peduzzi et al 2003). In the microoxic incubations, dark C-fixation activity of *Chr. okenii* increased proportionally with the amount of O_2_ added (Fig 3B&C) confirming that these bacteria obtained energy from aerobic respiration. The mean growth rate at an expected concentration of 15 μM O_2_ (0.082 ± 0.021 d^−^1) was about three times higher than at an expected concentration of 5 μM O_2_ (0.025 ± 0.008 d^−1^). Based on these growth rates and total cell abundances, we could estimate that *Chr. okenii* accounted for 31-42 % of the dark C-assimilation measured by bulk analyses (Fig. S3).

**Figure 3.**
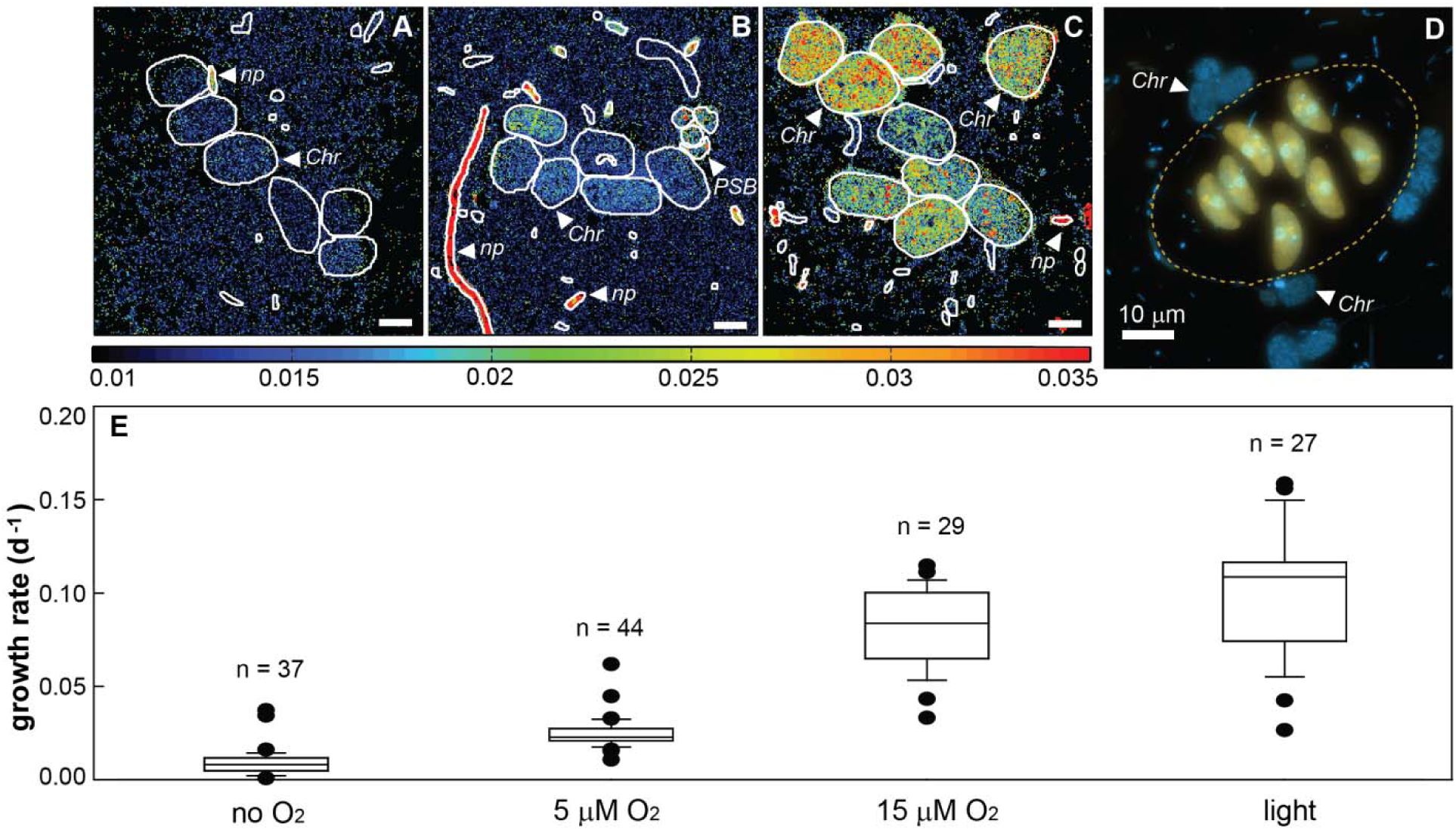
^13^CO_2_ uptake by individual *Chr. okenii* cells incubated in the dark with no O_2_, 5 μM (expected) dissolved O_2_, and 15 μM (expected) O_2_, and in the light with no added O_2_. Secondary ion images (A-C) show ratios of ^13^C/^12^C. Scale bars are 3 μm and bacterial cells are outlined in white. Epifluorescence microscopy image (D) of DAPI-stained *Chr. okenii* associated with a consortium of autofluorescent algal cells enclosed within an organic sheath (dotted orange line, defined based on ^12^C^14^N image in Fig S8). White triangles denote cells identified as: *Chr = Chr. okenii, PSB* = other purple sulfur bacteria, and *np* = non-phototrophic. (E) Box-and-whisker plots showing the range and the median growth rate of *Chr. okenii* under each incubation condition calculated from ^13^C-uptake per cell.

Clusters of cells reminiscent of the PSB *Thiodictyon* as well as a few other bacterial filaments and rod-shaped cells that were most likely aerobic, sulfide-oxidizing bacteria were also highly enriched in ^13^C (Fig 3B&C). This suggests that the capacity for aerobic respiration is taxonomically widespread in the anoxic chemocline.

Surprisingly, *Chr. okenii* growth rates in the light (0.100 ± 0.033) were only slightly higher than under microoxic conditions (Fig 3E) and their activity represented about 25 % of the total C-assimilation (Fig. S3). Since this incubation was performed over a full light-dark cycle, it is also possible that activity was restricted to the 13-h light period, which would yield a higher hourly growth rate. While it is difficult to disentangle phototrophic from chemoautotrophic carbon fixation in this case, the presence of abundant associations of *Chr. okenii* cells with eukaryotic algae (Fig. 3D) indicates that these PSB can also thrive off local light-driven O_2_ production. We could show that algae were photosynthetically active via the incorporation of ^13^CO_2_ into the eukaryotic cells (Fig. S10). Diffusion of O_2_ to attached *Chr. okenii* cells may have stimulated chemoautotrophic growth as they were also highly enriched in ^13^C.

*Chr. okenii* use energy from aerobic respiration not only for anabolic processes but for motility as well. We could confirm that *Chr. okenii* are highly motile both in the day and the night by performing dark field video microscopy (see Movie S1 in Supplementary Materials) of environmental samples obtained during the night and monitored in a dark room to avoid light-induced artefacts. Although the average night time swimming speed of *Chr. okenii* (9.9 μm s^−1^; see Fig. S11) was a third of the day time swimming speed (27 μm s^−1^; Sommer et al. 2017), it is clear that *Chr. okenii* remains motile even under dark conditions. This motility may in fact be key to bridging spatially separated gradients of oxygen and sulfide on which they grow.

### *Metagenomic insights into the* Chr. okenii *population in Lake Cadagno*

To assess the genomic potential for light-independent, aerobic sulfide oxidation by *Chr. okenii* in Lake Cadagno, we sequenced two metagenomes, one from the chemocline and one from a phototrophic, sulfide-oxidizing enrichment culture obtained from the lake (Table S1). From a combined metagenomics assembly, we reconstructed a high quality (90% complete, <1% contaminated) metagenome-assembled genome (MAG) of a PSB highly abundant in the sulfur-oxidizing enrichment culture (Fig. S12). The recovered MAG had a low average nucleotide identity ANI (<70%) to any sequenced *Chromatiaceae* genomes (data not shown). However, it encoded an rRNA operon, including a complete 16S rRNA gene with 99% sequence identity to the 16S rRNA gene of *Chr. okenii* (Imhoff *et al*., 1998; Tonolla *et al*., 1999), and thus likely represents a strain of *Chr. okenii* which is the type strain of the genus *Chromatium*. At this time, *Chr. okenii* has not been successfully isolated in pure culture, nor is there any published genome available for this organism.

The key metabolic process of *Chr. okenii* in Lake Cadagno is photoautotrophic sulfur oxidation. In accordance, the *Chr. okenii* MAG contained genes encoding for a sulfide : quinone reductase (*sqr*) and the full genomic inventory coding for a reverse-acting dissimilatory sulfite reductase (rDSR) pathway (Fig. 4). The operon structure of the rDSR encoding genes (*dsrABEFHCMKLJOPN*) was identical to the operon structure in the well described PSB model organism *Allochromatium vinosum* (Dahl *et al*., 2005), but no *dsrR* and *dsrS* gene were found. No genes encoding for sulfur oxidation via the SOX pathway, or homologues of sulfur globule proteins (*sgpABC*) typically found in PSB were detected in the draft genome. In line with its phototrophic metabolism, the *Chr. okenii* MAG showed the genomic potential for photosynthesis, with the genes encoding for a light harvesting complex 1 (*pufAB*) and a PSB-type photosynthetic reaction center (*pufLMC*) encoded in a single operon. Furthermore, the full genomic repertoire for a NADP-Me type C4 photosynthetic carbon assimilation cycle, and all genes (with exception of *cbbS* encoding for the small subunit of the ribulose-1,5-bisphosphate carboxylase/oxygenase) necessary for CO_2_ assimilation via the Calvin-Benson-Bassham (CBB) Cycle were present (Fig. 4).

**Figure 4.**
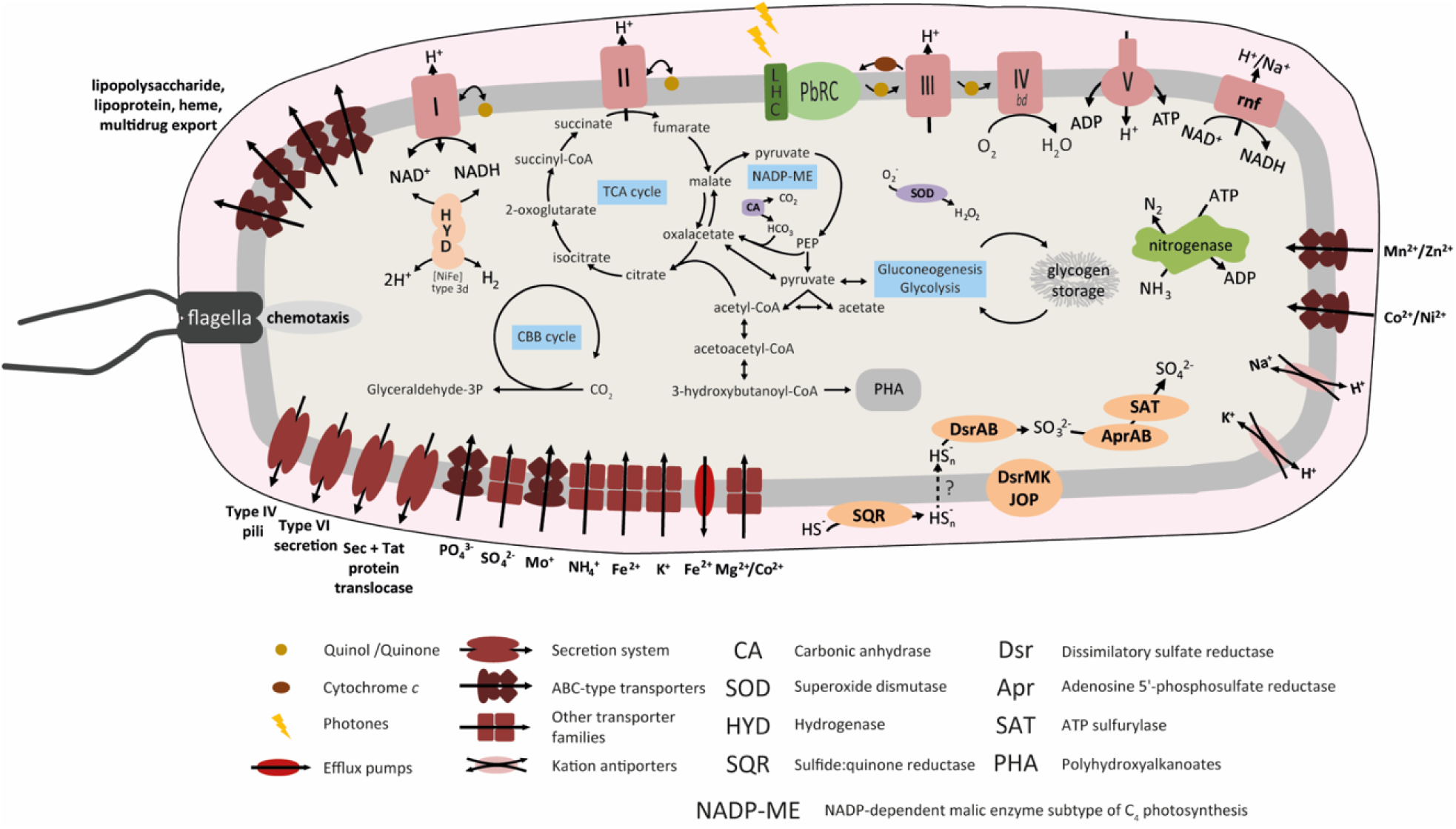
*Chr. okenii* cell illustration, showing the metabolic potential inferred from the metagenome-assembled genome with a particular focus on the genetic machinery implicated in photosynthesis, sulfur oxidation, aerobic metabolism, motility, glycogen and PHA storage, nitrogen fixation and transmembrane transport. The respiratory chain enzyme complexes are labeled with Roman numerals.

Many *Chromatiaceae* can grow chemoautotrophically, respiring oxygen under microoxic conditions (Kämpf and Pfennig, 1980). Cytochrome (Cyt) *c*-containing oxidases (e.g. Cyt *aa3*, Cyt *cbb3*) were not found in the *Chr. okenii* MAG. However, a Cyt *bd* type ubiqinol oxidase, known to function as sulfide-resistant O_2_-accepting oxidase in other *Gammaproteobacteria* (Forte *et al*., 2016), was identified (Fig. 4). Further, a plethora of genes related to heme *b* (*gltX, hemALBCD*, and *hemH*) and siroheme (*cysG*) synthesis, degradation (a heme oxygenase) and export (ABC-type heme exporter, *ccmABCD*), as well as hemerythrin-like metal binding proteins were encoded. Hemerythrin has been implicated in binding of oxygen for delivery to oxygen- requiring enzymes, for detoxification, or for oxygen sensing in motile, microaerobic prokaryotes (French *et al*., 2007). The presence of these oxygen-dependent enzymes, as well as a key oxidative stress defense enzyme superoxide dismutase (SOD), support the idea that *Chr. okenii* may be facultatively microaerobic. A complete set of genes for flagellar biosynthesis (*fliDEGHJKLMNOPQRW*, *flgABCDEFGHIK, flhAB*) and flagellar motor proteins (*motAB*) confer motility to this bacterium.

Several other genes revealed interesting metabolic capacities of *Chr. okenii*. A cytosolic bidirectional [NiFe] type 3d hydrogenase and a nitrogenase were encoded in the MAG (Fig. 4), implicating the potential for involvement of *Chr. okenii* in nitrogen fixation and hydrogen oxidation which has previously been overlooked. Additionally, the *Chr. okenii* MAG encoded a glycogen synthase and a glycogen debranching enzyme, as well as the full genomic repertoire necessary for polyhydroxyalkanoate (PHA) biosynthesis. This is consistent with the detection of glycogen in our biogeochemical profiles of the chemocline. Finally, it is possible that novel terminal oxidases are among the hypothetical genes that could not be assigned any known function.

## CONCLUSIONS

It is intriguing that oxygen should play a major role in sulfide oxidation in the ostensibly anoxic chemocline of Lake Cadagno, especially by purple sulfur bacteria generally thought to lead an anaerobic lifestyle. To explain the coupling of oxygen and sulfide consumption in the oxygen- and sulfide-free chemocline of Lake Cadagno, we sketched a diagram of the transport processes likely driving biological activity there (Fig. 5). As described in Sommer *et al*., (2017), active convection of the chemocline can be driven by the formation of sinking bacterial plumes. Combined with turbulence induced by the breaking of internal waves at the sides of the lake basin, these convective currents may entrain sulfide and oxygen at the boundaries of the chemocline and fuel populations of sulfide-oxidizing *Chr. okenii* there.

**Figure 5.**
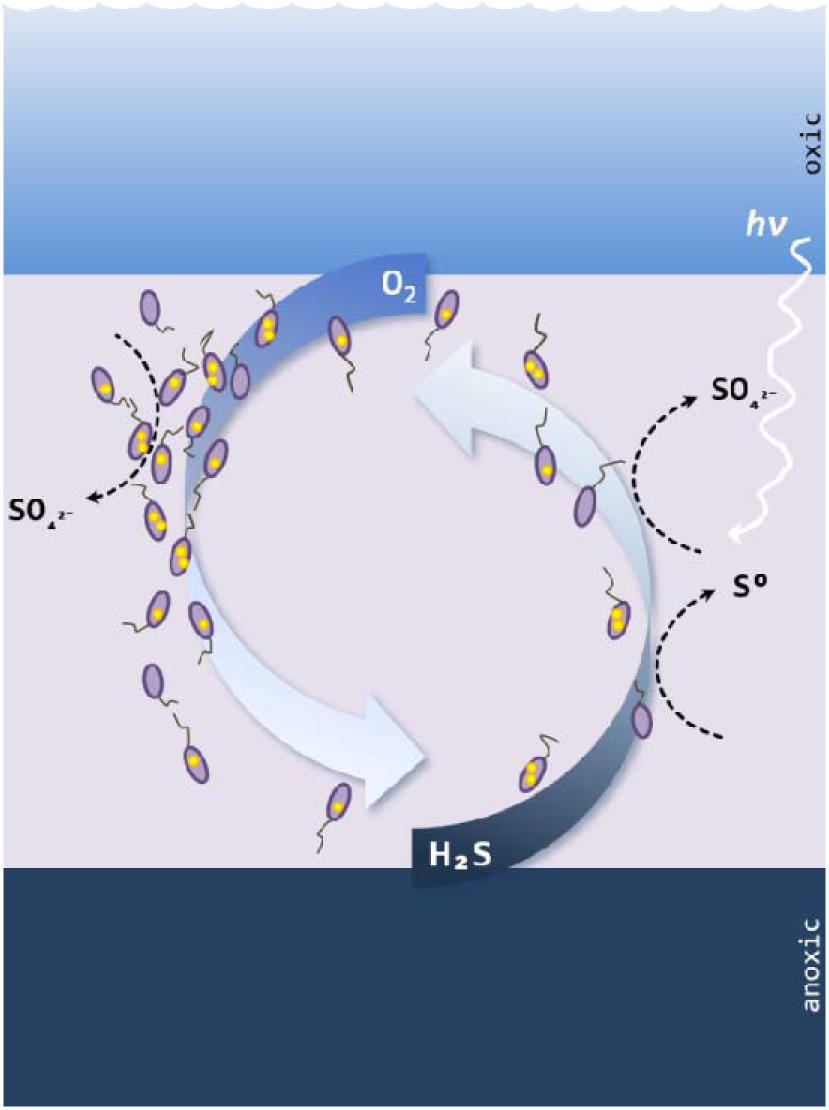
Schematic of phototrophic and aerobic sulfide oxidation processes in the Lake Cadagno chemocline. Convection in the chemocline may be driven by a combination of turbulence and sinking bacterial plumes, represented by the large number of descending *Chr. okenii* cells on the left. As a result, oxygen and sulfide are entrained into the chemocline and immediately consumed by purple sulfur bacteria, keeping concentrations of these compounds below detection limits. *Chr. okenii* cells, depicted with internal sulfur globules (yellow dots), are pulled in the direction of their flagellar bundle.

Sulfur-oxidizing bacteria have previously been reported to bridge distances between pools of electron donors and acceptors by intracellularly storing and transporting S^0^ and NO_3_^−^ between redox zones (Fossing *et al*., 1995; Jørgensen and Gallardo, 1999) and even by transferring electrons along nanowires (Pfeffer *et al*., 2012), but aerobic sulfide oxidation in Lake Cadagno represents a new mechanism of electron acceptor/donor coupling across large distances. After entrainment into the chemocline, dissolved oxygen and sulfide are consumed so rapidly that they remain below detection limits. The physical and biological processes described here may therefore provide clues to sulfide oxidation in other anoxic environments such as the Black Sea where the mechanism of sulfide removal is not completely understood. Clearly, the biochemical limits to oxygen utilization are far below current definitions of anoxia and demonstrate that aerobic respiration is possible in so-called “anoxic” lacustrine (Milucka *et al*., 2015) and marine (Garcia-Robledo *et al*., 2017) waters.

Although purple sulfur bacteria are generally considered anaerobes, we propose that *Chr. okenii* in the Lake Cadagno chemocline aerobically respire upwards-diffusing sulfide in addition to performing phototrophic sulfide oxidation as previously reported (e.g. Tonolla et al. 1999; Musat et al 2008; Storelli et al. 2013). Our direct evidence for oxygen-driven C-fixation and continued motility of *Chr. okenii* in darkness is also supported by the presence of a potential high-affinity oxidase in the metagenome-assembled genome. In contrast to observations from laboratory cultures, *Chr. okenii* may have a very different metabolism in the environment where high fluxes of substrates rather than absolute substrate concentrations fuel microbial activity.

Our nanoSIMS measurements revealed that in addition to *Chr. okenii*, a number of phototrophic and non-phototrophic cells are also highly active under aerobic, sulfide-oxidizing conditions. Based on the diversity of genes encoding for high-O_2_ affinity terminal oxidases found in our Lake Cadagno metagenome, it appears that some other purple sulfur bacteria as well as a handful of *Gamma-* and *Epsilonproteobacteria* might aerobically oxidize sulfide (Fig. S13). Interestingly, two species of strictly anaerobic green sulfur bacteria from Lake Cadagno also possessed genes encoding for terminal oxidases, though further study will be required to elucidate the role of these different bacterial populations in chemotrophic sulfide oxidation.

## EXPERIMENTAL PROCEDURES

### Sampling

The meromictic Lake Cadagno is situated in the Piora Valley in the Swiss Alps at an altitude of 1921 m. Data presented here were collected during field campaigns in September 2013, August 2014, June 2015 August 2015, and August 2018. In 2013 and 2014 *in situ* measurements were performed with a profiling ion analyzer (PIA; see Kirf *et al*., 2014 for description) lowered from a platform anchored at the deepest part of the lake (20.7 m). Conductivity, turbidity, depth (pressure), temperature and pH were measured with a multi-parameter probe (XRX 620, RBR). Dissolved oxygen was recorded online with a type PSt1 normal (detection limit 125 nM) micro- optode and a type TOS7 trace (reliable detection limit 50-100 nM) micro-optode (PreSens). The oxygen sensors were calibrated by parallel Winkler titrations. Water samples for chemical analyses and cell counts were collected with a rosette syringe sampler equipped with twelve 60-ml syringes triggered online at selected depths. Due to a technical failure of the PIA, the 6 AM profile in August 2014 and all subsequent profiles were measured with an SBE 19 plus V2 CTD probe (Sea-Bird Electronics) equipped with sensors for pressure, temperature and conductivity, and with additional sensors for turbidity (WET Labs Eco), oxygen (SBE 43), pH (18-1) and two fluorescence wavelengths (WET Labs ECO-AFL). The detection limit of the SBE 43 oxygen probe was about 1 μmol/l. In parallel with *in situ* measurements, water for chemical analyses was pumped to the surface through Neoprene tubing attached to the CTD and filled into 60-ml syringes (rinsed 2 X with *in situ* water) on board. Two parallel metal plates of diameter ~15 cm attached to the submersed end of the tubing served to channel water horizontally, resulting in more discrete vertical profiling.

Water samples in syringes were aliquoted on board immediately after collection. Samples for sulfate analyses were filtered (0.22 μm pore size) directly into sterile Eppendorf vials. Sulfide samples were fixed with Zn-acetate to a final concentration of 0.1 % (w/v). Biomass was concentrated onto GF/F filters (Whatman) and stored at −20°C for analyses of intracellularly stored elemental sulfur and organic carbon compounds. Filtrate (0.22 μm pore size) was also collected and frozen at −20°C for metabolome analysis of dissolved compounds. Water for dissolved inorganic carbon (DIC) analysis was filled into 12-ml Exetainers (Labco Limited) and capped before injecting 100 μl of saturated HgCl_2_ solution to terminate microbial activity. Biomass samples for determination of background ^13^C in the particulate organic carbon (POC) were concentrated onto pre-combusted (6 h at 600 °C) GF/F filters. Samples for DNA analysis were collected from the chemocline in August 2014 by concentrating microbial cells on 0.22 μm GTTP-type polycarbonate filters (Millipore) on site and freezing at −20°C until further processing. Additional water for cultivation and motility experiments was pumped directly from the chemocline into 1-L Duran bottles or 250-ml serum bottles and sealed with butyl rubber stoppers without a headspace to maintain anoxic conditions.

### Chemical Analyses

Sulfide was measured using the colorimetric method of Cline (1969). Sulfate was measured on a 761 Compact ion chromatograph (Metrohm) equipped with a Metrosep A SUPP 5 column. Intracellular sulfur on filters was extracted by sonication in methanol for 15 min in an ice bath. Samples were analyzed on an Acquity H-Class UPLC system (Waters Corporation) with an Acquity UPLC BEH C18 column coupled to a photodiode array (PDA) detector using UPLC-grade methanol as eluent. Data was acquired and processed using the Empower III software.

Intracellular glycogen was analyzed following the procedures of the assay kit (MAK016, Sigma Aldrich). Briefly, cells were extracted by scraping them from GF/F filters and homogenizing in 200 μL extraction buffer and centrifuged two times to clear the supernatant. The supernatant was analyzed fluorometrically after incubation with enzyme mix and fluorescent peroxidase substrate. Intracellular PHA was analyzed using the protocol from Braunegg *et al*., (1978). Hydrolyzation of the polymer and conversion to a methyl-ester of the monomeric hydroxyalkanoate fraction was done in acidified alcohol solution (6% H_2_SO_4_ in methanol) and chloroform under heating (100°C, 2 h). After addition of water and phase separation the organic phase was analyzed with GC-MS (Agilent 7890B GC connected to Agilent 5977A MSD) to detect the methylhydroxyalkanoates using the following settings: Agilent 30 m DB-5-MS column, splitless injection of 1 μl, temperature program was 50°C for 1 min than heating 10°C/min until 120°C followed by 45°C/min until 320°C and hold for 5 minutes. Benzoic acid was used as internal standard in each sample and quantification was done with pure polyhydroxybutyrate standard (Sigma Aldrich).

Sulfate reduction rates were measured by adding the radiotracer ^35^SO_4_^2−^ (5 MBq) to anoxic lake water in 50-ml glass syringes and incubated in the dark. A solution of unlabeled Na_2_S was added to a final concentration of ~50 μmol·l^−1^ as a background sulfide pool in case of sulfide re-oxidation. At each sampling point, 10 ml of sample was dispensed into 5 ml of 20% (w/v) Zn-acetate. Reduced sulfur species (e.g. sulfur and sulfide as ZnS) were separated out via the chromium distillation method described in Kallmeyer *et al*., (2004) and the radioactivity per sample was determined via scintillation counting (Packard 2500 TR).

### Confocal Raman spectroscopy

In a glove box under 90:10 N_2_-CO_2_ atmosphere, a drop of fresh sample from the chemocline was mounted between two glass coverslips and sealed with electrical tape to prevent contact with air. A polysulfide solution containing 5.06 g Na_2_S · 9H_2_O and 5.8 g elemental sulfur per 100 ml H_2_O, with a final pH of 9.5 and sulfide concentration of 210 mM, was used as reference. Measurements were conducted with an NTEGRA Spectra confocal spectrometer (NT-MDT) coupled to an inverted Olympus IX71 microscope. The excitation light from a 532-nm solid-state laser was focused on the sample through an Olympus 100X (numerical aperture [NA], 1.3) oil immersion objective. Raman scattered light was collected by an electron-multiplying charge-coupled device (EMCCD) camera (Andor Technology) cooled to −70°C. Spectra were recorded between 0 and 4,500 cm^−1^ with a spectral resolution of 0.2 cm^−1^ and analyzed with the NT-MDT software Nova_Px 3.1.0.0.

### Flux Calculations

Turbulent fluxes (*J*) of sulfide, sulfur, sulfate, and oxygen at the chemocline were calculated assuming steady state by applying Fick’s first law: *J=*−*D∂C/∂x*. For sulfide, sulfate, and oxygen we used the turbulent diffusion coefficient (D) of 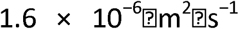 from (Wüest, 1994) corresponding to turbulence at the Lake Cadagno chemocline boundaries. For sulfur gradients within the well-mixed chemocline the coefficient 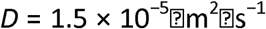 from Wüest (1994) was used. The change in concentration (*∂C*) was computed for each species over the depths with the steepest gradients. Oxygen and sulfide fluxes were determined for the regions immediately above and below the chemocline, defined as the zone of constant conductivity.

### Potential oxygen consumption rate measurements

Potential O_2_ consumption rate measurements were conducted as described in Holtappels et al. (2014). Incubations were performed in 250 ml Winkler bottles using trace oxygen sensor spots (Ø5mm, TROXSP5, Pyroscience) glued to the inner glass wall and read from the outside using a 4-channel fiber-optic oxygen meter (FireStingO2, Pyroscience) recording O_2_ concentration every 10 seconds. The glass stopper of the Winkler bottle was pierced with a glass tube (11 cm long, inner diameter 1.5 mm), to allow for pressure compensation due to temperature-induced volume changes and to facilitate the addition of small amounts of reagents. Oxygen transport through this opening was limited by diffusion through a narrow passage over a long distance. Tests with degassed sterile water showed negligible O_2_ concentration changes of 3-25 nM·d^−1^ (figures available on request). Water inside the incubation bottles was gently mixed by small glass-coated magnets driven by an adjacent magnetic stirrer. The incubation bottles were placed in a temperature-controlled water bath set to 4.8°C (+/− 0.1°C). A two-point calibration of the trace optode spots was performed by using degassed water to which a known amount of O_2_-saturated water was added through the glass tube.

Anoxic chemocline water was pumped onboard a sampling platform and distributed directly into incubation vessels via Neoprene tubing, avoiding atmospheric contamination as described above. Bottles were placed immediately into the water bath and incubated in the dark for 18 hours. Initial O_2_ values were between 6 and 8 μM. Over a period of 12 hours, H_2_S was added 5 times by injecting up to 300 μl of a 4 mM Na_2_S stock solution to a final concentration of 1.6 μM. Between 4-6 ml O_2_-saturated lake water from the same depth was periodically added to maintain O_2_ concentrations between 5 and 11 μM. O_2_ consumption rates were calculated from linear regressions over 10-30 min time intervals.

### Stable isotope incubations

250-ml serum bottles containing Lake Cadagno chemocline water collected in August 2018 were bubbled for 10 min with N_2_ to remove any contaminant oxygen. Four different incubation conditions were set up to test microbial carbon fixation under the following conditions: no O_2_ + dark, no O_2_ + light, medium O_2_ + dark, and high O_2_ + dark. To each of the dark incubations, 50 ml of water was removed and replaced with N_2_ gas. Solutions of Na_2_S and ^13^C-DIC were added to final concentrations of 25 μM and 1 mM (36-39 atom %), respectively, except for the light incubation which received 200 μM (8 atom %) ^13^C-DIC. Volumes of 0.918 ml and 2.8 ml air were added to the dark, aerobic incubations to achieve headspace O_2_ concentrations of 129 and 388 μM, respectively, resulting in expected dissolved O_2_ concentrations of 5 and 15 μM, respectively, at the start of the incubations (Garcia and Gordon, 1992). All bottles were gently shaken for several minutes to equilibrate the water and headspace and then placed in a 10°C water bath in the dark for 18 h, except for the light condition which was incubated separately under natural light, 0.1 – 2 % incident daylight, for a full light-dark cycle. Equilibration with the headspace was enhanced throughout the incubation by stirring with glass-coated magnetic stir-bars at minimum speed.

In order to verify the consumption of oxygen, two bottles prepared without ^13^C-DIC and injected with 0.918 ml and 2.8 ml of air were incubated in parallel with needle optodes (FirestingO2, Pyroscience) immersed in the liquid phase. Dissolved O_2_ concentrations were recorded every 3 min and are shown in Fig. S9.

At the end of the experiment, bottles were sacrificed for analysis of bulk and single-cell ^13^C uptake. Water for ^13^C-DIC and ^13^C-POC was sampled as described above. For nanoSIMS, water samples were fixed with formaldehyde (2 % v/v) overnight at 4°C before filtering onto 0.2 μm Au/Pd-sputtered polycarbonate filters (Millipore) and stored at −20°C until processing.

### Mass spectrometry isotope uptake analysis

The labeling % of ^13^C-DIC was determined by diluting Hg-killed water 1:10 in MilliQ and transferring 3 ml into a 12 ml Exetainer with an N_2_ headspace. Samples were then acidified with the addition of ~100 μl concentrated H_3_PO_4_ so the outgassed ^13^CO_2_ could be analyzed by cavity ring-down spectroscopy (G2201-i Picarro).

The incorporation of ^13^C from ^13^CO_2_ into biomass was measured by combustion of the particulate organic carbon fraction on GF/F filters. Filters were decalcified by incubation with 37% fuming HCI overnight, dried in an oven at 60°C, and then packed into tin capsules for combustion analysis. The C isotopic composition was determined on an automated elemental analyzer (Thermo Flash EA, 1112 Series) coupled to a Delta Plus XP IRMS (Thermo Finnigan).

### Nanometer-scale secondary ion mass-spectrometry

Circular pieces (5-mm diameter) were punched out of the Au/Pd-sputtered filter and regions of interest were marked with a laser micro-dissection microscope (LMD) (DM 6000 B, Leica Microsystems) before analysis with the nanoSIMS 50L (Cameca) at the Max Planck Institute for Marine Microbiology in Bremen, Germany. For each field of view, the sample was pre-sputtered with a primary Cs^+^ beam of 300 pA, and then measured by rastering a primary Cs^+^ beam with a diameter of < 100 nm and a beam current between 1.0 and 1.5 pA over the area. Secondary ion images of 12C^−^, 13C^−^, 12C14N^−^, 31P^−^ and 32S^−^ were recorded simultaneously using 5 detectors together with the secondary electron (SE) image ranging from 10 × 10 μm to 50 × 50 μm in size and corresponding to an image size of 256 × 256 and 512 × 512 pixels, respectively, using a dwell time of 1 ms per pixel. Up to 60 planes were recorded for each imaged area.

Images were processed using the Matlab-based Look@NanoSIMS software (Polerecky *et al*., 2012). Regions of interest (ROIs) were drawn around cells based on the 12C14N^−^ signal overlain on SE images and *Chr. okenii* cells were identified based on morphology, size, and pigmentation in parallel light-microscope images. Cellular ^13^C atm. % was calculated from ^13^C/^12^C values of individual ROIs. The background (cell-free polycarbonate filter) ^13^C content was also evaluated in every field of view for comparison. Growth rates were calculated assuming a linear growth rate where one cell division results in a maximum of ½ the labeling percentage:

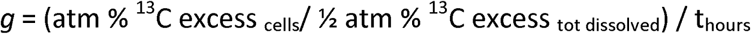

Cellular C-uptakes rates were calculated based on the relationship between biovolume and C-content from Verity *et al*., (1992), assuming an average biovolume of spherocylinders cells of 65 μm^3^ based on direct measurements of 70 *Chr. okenii* cells.

### Motility analysis

Water samples containing *Chr. okenii* cells were collected under anoxic conditions from the chemocline during the night, protected from artificial light with aluminum foil, and analyzed immediately on site. Motile cells were transferred via a degassed glass syringe to a sealed rectangular millimetric chamber (dimensions 20 mm × 10 mm × 2 mm) prepared using glass slides separated by a 2-mm thick spacer, which provided an anoxic environment during motility characterization. Experiments were conducted in a dark room, and imaging was performed using the dark field microscopy mode at 25 fps, with the lowest intensity illumination. No transient response was observed right at the start of the imaging, and the swimming velocity remained steady throughout the duration of the measurements. This is in contrast to swimming behavior at higher light intensities where the swimming cells exhibited a positive phototactic response (Sommer *et al*., 2017). We could therefore rule out a light-induced effect on motility at the minimum illumination level used for our measurements. Videos of swimming cells were acquired and subsequently analyzed using the ImageJ Particle Tracker routine to obtain the coordinates of the cells (geometric centers) at each time interval. These were used to calculate the swimming speeds and extract the trajectories of individual cells.

### DNA extraction, sequencing, and analysis

Two metagenomes were sequenced: one from the chemocline and one from a phototrophic, sulfide-oxidizing enrichment from the lake. The enrichment culture was obtained by preparing agar-stabilized sulfide-gradient tubes as described in Schwedt et al. (2012) using sterile-filtered Lake Cadagno water amended with vitamins and trace elements as described for cultivation of purple sulfur bacteria (Eichler and Pfennig, 1988). After inoculation with water from the chemocline, cultures were incubated for approximately 3 months in indirect light at 15°C until development of a deep purple color indicative of growth of PSB.

DNA was extracted from polycarbonate filters with the Ultra Clean MoBio PowerSoil DNA kit (MoBio Laboratories, Carlsbad, USA) according to the manufacturer’s protocol with the following modification: the bead beating step was reduced to 30 sec followed by incubation on ice for 30 sec, repeated 4x. The DNA was gel-purified using SYBR Green I Nucleic Acid Gel Stain (Invitrogen) and the QIAquick Gel Extraction Kit (Qiagen) according to the accompanying protocols. DNA concentration was determined fluorometrically at 260 nm, using the Qubit 2.0 Fluorometer and the Qubit dsDNA HS Assay KIT (Invitrogen) and sent to the Max Planck-Genome Centre (Cologne, Germany) for sequencing. The metagenome was sequenced (100 bp paired end reads) by Illumina HiSeq (Illumina Inc., USA) sequencing following a TruSeq library preparation. Metagenomic reads were adapter- and quality-trimmed (phred score 15, bbduk function of the BBMap package, https://sourceforge.net/projects/bbmap/) and paired-end reads were *de novo* assembled with the uneven depth assembler IDBA-UD (Peng *et al*., 2012).

The metagenome assembly was binned based on tetranucleotide frequencies, differential coverage, taxonomic classification, and conserved single-copy gene profiles with the Metawatt binning software (version 3.5.2; Strous *et al*., 2012). The completeness and contamination of the binned MAGs was evaluated with CheckM (Parks et al 2014). The bulk metagenome and the MAG identified as *Chr. okenii* were automatically annotated in IMG (Markowitz et al 2011), and the *Chr. okenii* MAG was manually screened for the presence of genes of interest to this study. Assembled data is available in IMG, under the IMG genome IDs 3300010965 (bulk assembly) and 2700988602 (*Chr. okenii* MAG).

Cytochrome *c* and quinol oxidase subunit I and the cytochrome *bd* oxidase subunit I proteins were extracted from the metagenome via Pfam motif search (PF00115 and PF01654, respectively), and manually inspected to exclude NO reductase subunit I (NorB) proteins and proteins shorter than 100 aa prior to downstream analysis. A total of 130 out of 178 identified cytochrome *c* and quinol oxidase subunit I proteins and 76 out of 82 recovered cytochrome *bd* oxidase subunit I proteins passed the manual curation step. For phylogenetic tree reconstruction, the manually vetted protein sequences and a selection of reference sequences recovered from NCBI were aligned using MAFFT (Katoh *et al*., 2002) and IQ-TREE was used to construct the phylogenetic tree with automated best model prediction and a total of 1000 bootstrap replicates (Trifinopoulos *et al*., 2016; Hoang *et al*., 2017). Thereafter, a selection of sequences closely related to known sulfur oxidizing microorganisms alongside the respective reference sequences were used for reconstruction of the phylogenetic trees presented in Fig. S11. The output files were uploaded to iTOL for visualization (Letunic *et al*., 2016).

## ACKNOWLEDGEMENTS

We are grateful to the 2014 and 2015 Cadagno Field Expedition Teams from EAWAG and MPI Bremen for assistance in the field, and to the Alpine Biology Center Foundation (Switzerland) for use of its research facilities. We would especially like to thank Abiel Kidane and Sten Littman for help with nanoSIMS measurements as well as Nadine Rujanski, Gabriele Klockgether, and Gaute Lavik for mass spectrometry analyses. Funding was provided by the International Max Planck Research School of Marine Microbiology, the Max Planck Society, and the Deutsche Forschungsgemeinschaft (through the MARUM Center for Marine Environmental Sciences). A.S. was supported by the Human Frontier Science Program (Cross Disciplinary Fellowship, LT000993/2014-C).

