## Supplementary Figures for "Dark aerobic sulfide oxidation by anoxygenic phototrophs in anoxic waters"

Figure S1| Profiles of the Lake Cadagno water column from 2015 taken every ~4 hrs for 48 hrs. Sunset was at 20:53 and sunrise at 6:11. Some sulfide accumulates in the chemocline in the dark and presumably reacts with $S^0$ to form polysulfides which is detected as sulfide by the Cline method. $S^0$ was measured as particulate (intra- and extracellular) sulfur on 0.7 um filters and included polysulfides which would have spontaneously oxidized during filtration. Oxygen data was not available for profiles 1 and 8.

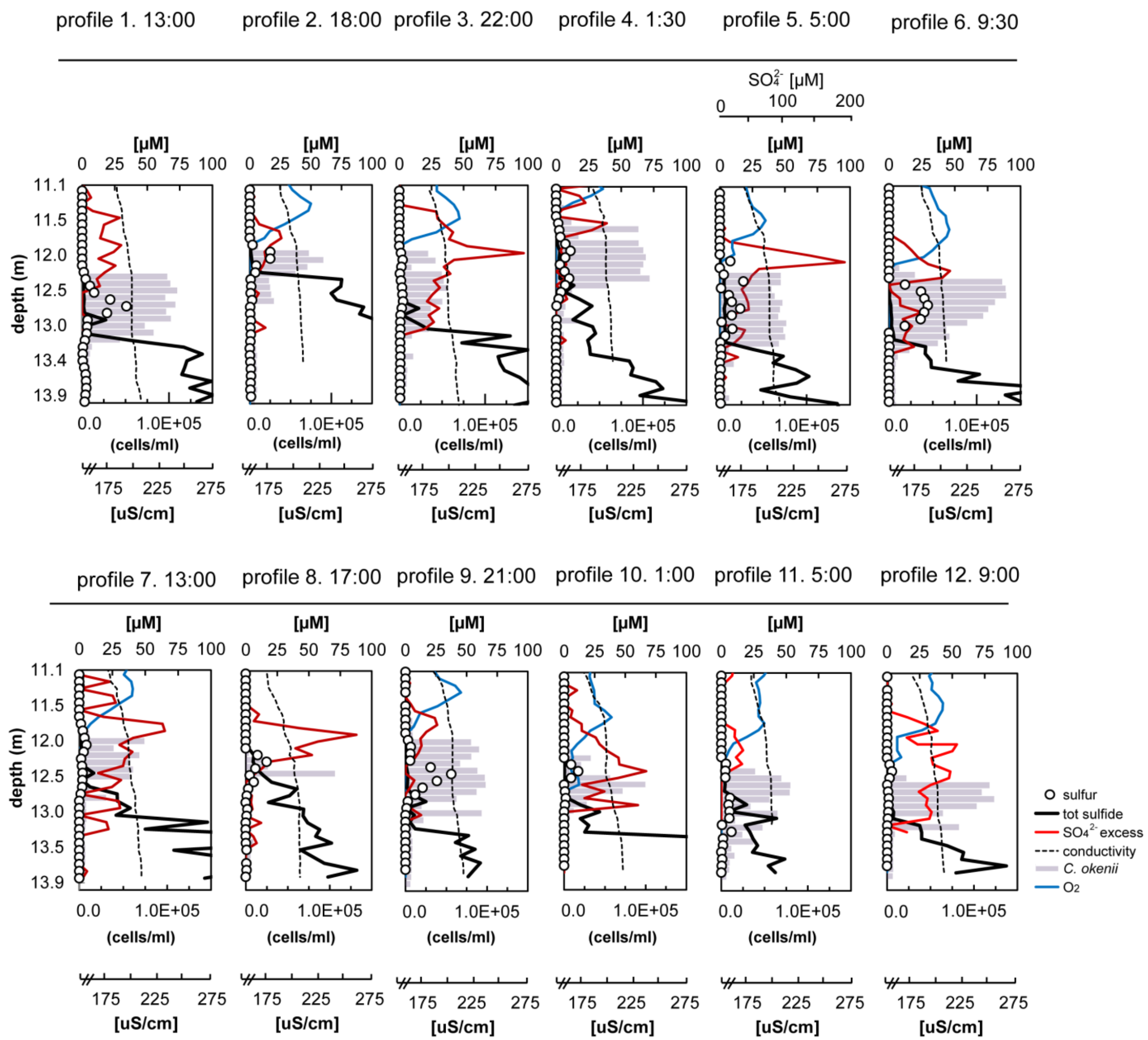

Figure S2: Profiles of the Lake Cadagno water column in Sept 2013 and Aug 2014 showing gradients of sulfide and oxygen, maximum turbidity in Nephelometric Turbidity Units (NTU), and temperature-conductivity ratios (T/C) in arbitrary units (a.u.). For 2014 profiles photosynthetically available radiation (PAR) and biogenic sulfate are also shown. September 2013 profiles were taken at midday (12:00), sunset (19:00) and after 1 h (20:00) and 4 h (23:00) of darkness. August 2014 profiles were taken at the end of the day (19:00) and just before sunrise after a full period of darkness (6:00).

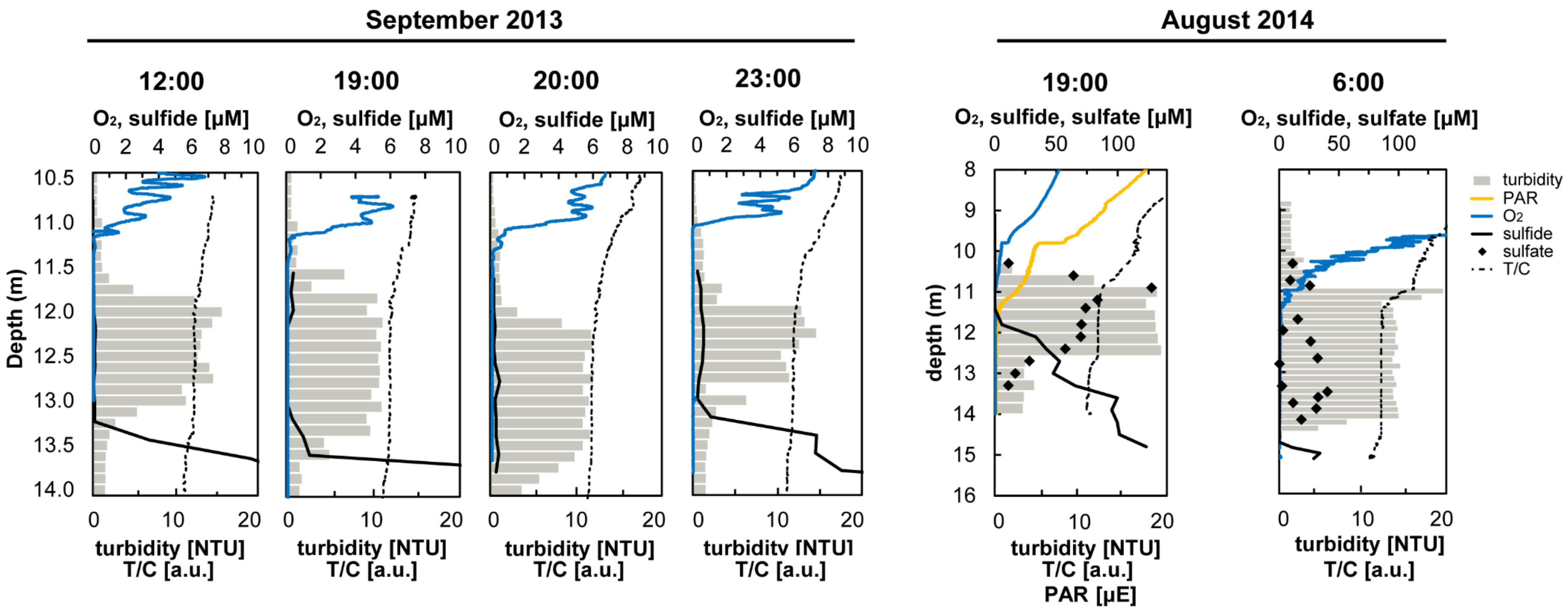

Figure S3| C-assimilation rates of *Chr. okenii* calculated from nanoSIMS measurements of 13C/12C ratios in single-cells compared to bulk C-assimilation rates measured in anoxic, low microoxic (5 µM expected $O_2$), medium microoxic (15 µM expected $O_2$), and light incubations performed in August 2018.

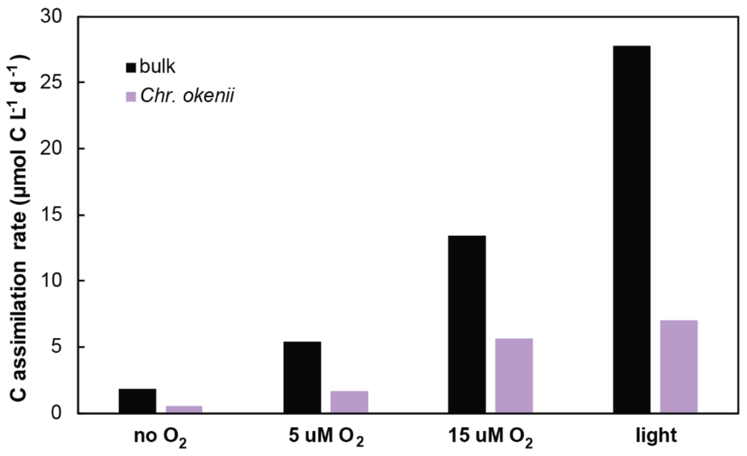

Figure S4| (A) Light microscope image of *Chr. okenii* cells with intracellular sulfur inclusions. Scale bar is 10 µm. (B) Raman spectrum of a sulfur inclusion from a living *Chr. okenii* cell in an environmental sample (upper spectrum) compared to a polysulfide standard (middle spectrum) and a cyclooctasulfur standard (lower spectrum). Exposure time was 0.5 sec.

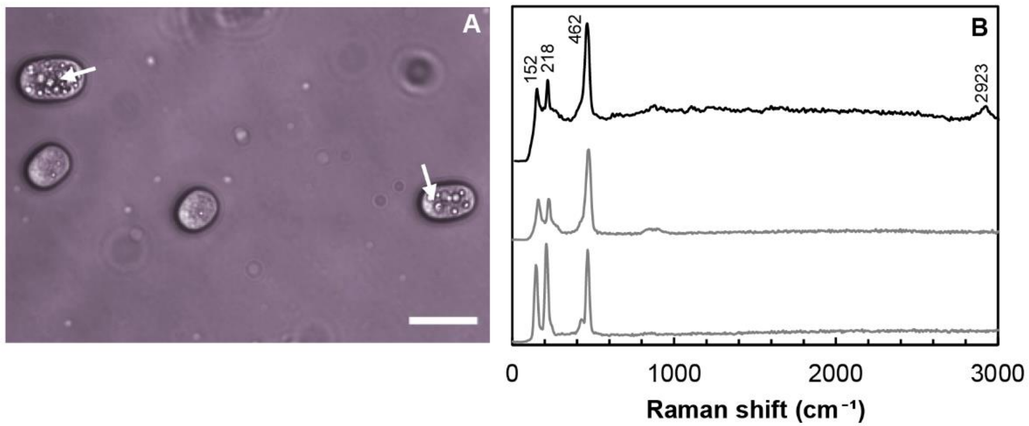

Figure S5| The day-night dynamics of various sulfur compounds in the Lake Cadagno chemocline plotted over 48 hours: (A) total $S^0$ integrated over the depth of the chemocline, (B) the total sulfide flux, (C) residual sulfide accumulated in the chemocline and (D) the total (upwards and downwards) flux of biogenic sulfate. Shaded regions represent dark periods.

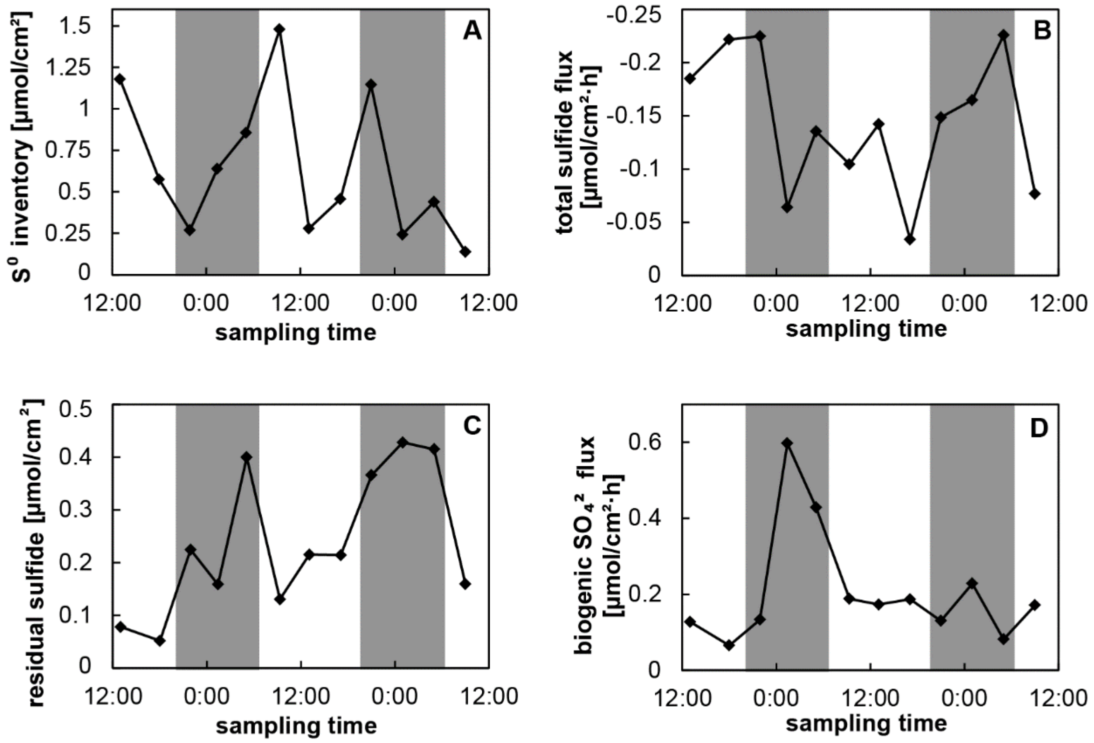

Figure S6| Glycogen concentration in the chemocline is plotted in picograms per *Chr. okenii* cell for one day (13:00) and one night (1:30) profile from 2015.

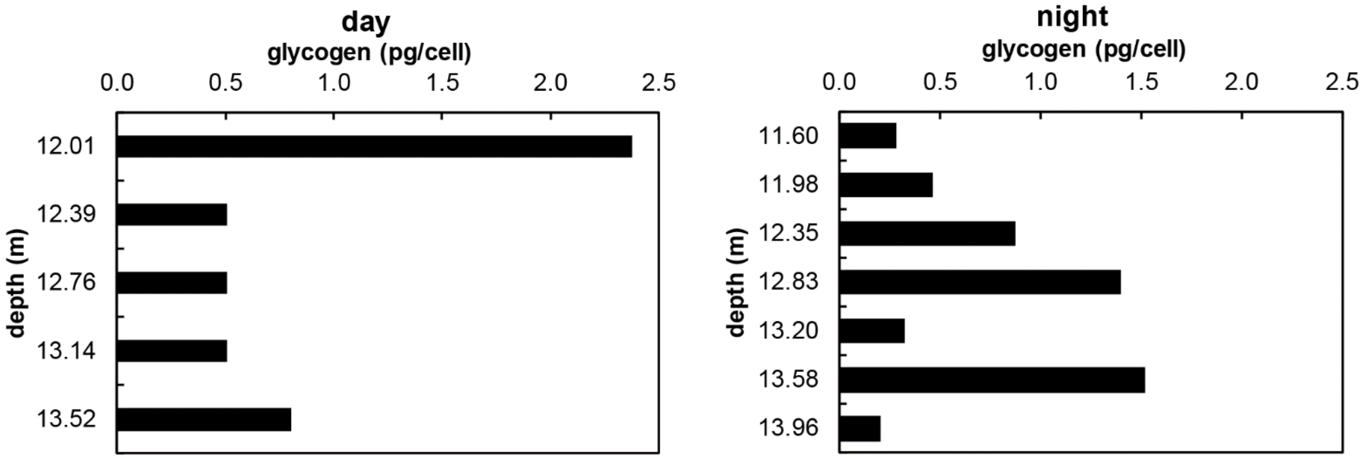

Figure S7| Examples of plots of sulfate concentration versus conductivity values from Lake Cadagno profiles measured in Aug 2014 at midday (16:00), sunset (19:00), and just before sunrise (6:00). A 1:1 mixing line was drawn between the end-member values from just above and below the chemocline. Points falling above this sulfate-conductivity mixing line represent excess sulfate.

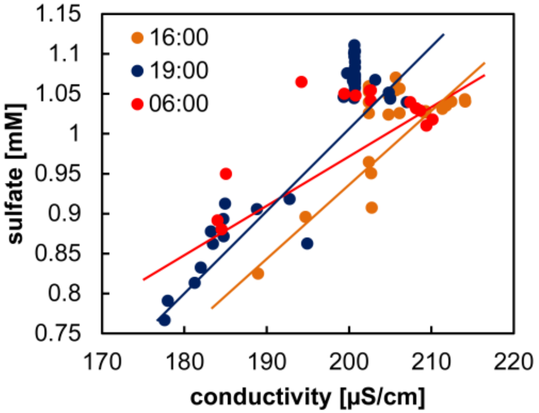

Fig S8| (A) $O_2$ concentration in $O_2$ consumption experiments were measured every 10 sec for 18 hours. Additions of $H_2S$ are marked by grey arrows. $O_2$ consumption rates 'before' and 'after' $H_2S$ consumption are calculated along the orange marked intervals. Red intervals demark rates during $H_2S$ consumption. (B) Summary of $O_2$ consumption rates before, during and after the $H_2S$ consumption. The background rates during the $H_2S$ experiment were slightly increased when compared to the rates at the very beginning and at the very end of the incubation. (C) Mean and standard deviation of all measured $O_2$ consumption rates before, during and after $H_2S$ consumption.

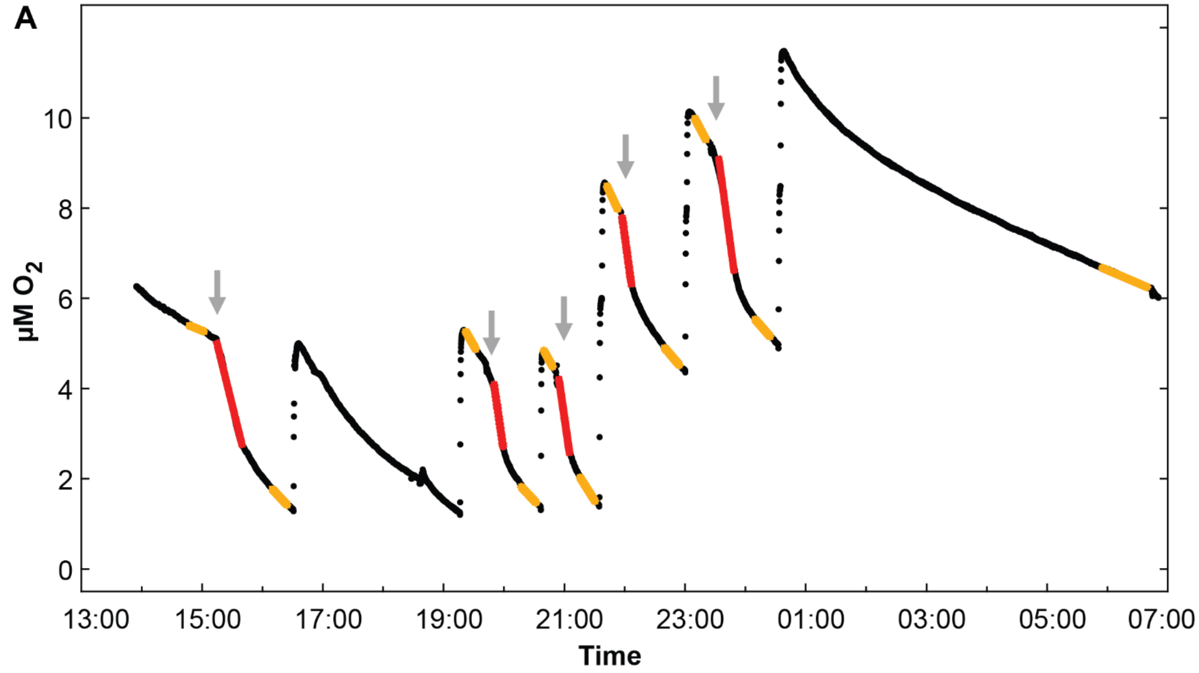


B

| $Na_2S$ addition (μM) | Before (μM/h) | During (μM/h) | After (μM/h) |
|---|---|---|---|
| 2.5 | -0.55 | -5.42 | -1.43 |
| 2.5 | -2.32 | -9.21 | -1.42 |
| 2.5 | -2.41 | -9.30 | -2.13 |
| 2.5 | -2.87 | -9.84 | -1.56 |
| 5 | -2.53 | -9.89 | -1.53 |
| mean | -2.14 | -8.73 | -1.61 |
| std. dev. | 0.81 | 1.68 | 0.26 |

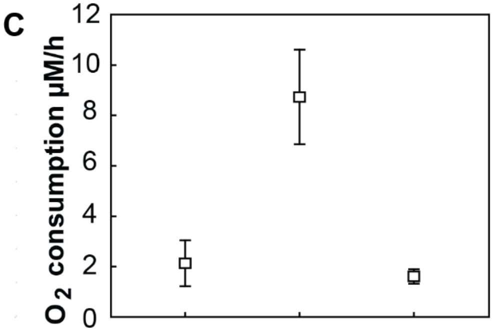

Fig S9| $O_2$ concentrations in bottles incubated in parallel to $^{13}C$-isotope labeling incubations performed in August 2018. The low $O_2$ and medium $O_2$ incubations were injected with air to expected dissolved $O_2$ concentrations of 5 µM and 15 µM $O_2$, respectively, at the start of the incubations. Consumption rates in the first hour were 8 µM/h which is the same as measured experimentally in August 2013 (Fig S8).

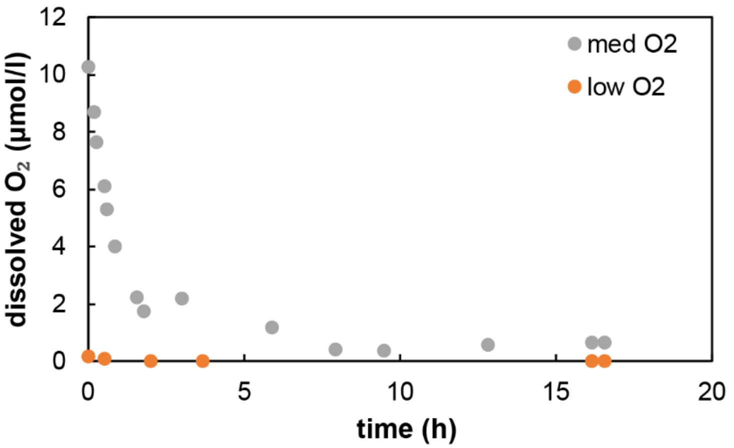

Figure S10| Secondary-ion images of *Chr. okenii* cells associated with a consortium of eukaryotic algal cells incubated with $^{13}C$-DIC in the light. The $^{13}C/^{12}C$ ratio (left) and $^{12}C^{14}N$ abundance (center) reveals $^{13}C$ uptake into *Chr. okenii* and algal cells but not the organic sheath visible in the parallel scanning electron image (right). Note that cell appear slightly less enriched than Fig 3C due to the lower amount of $^{13}C$-bicarbonate added (7 atom %). Cells imaged by epifluorescence microscopy (Fig 3D) were displaced during removal of the coverslip and embedding material prior to nanoSIMS imaging. Cell outlines are shown in white and *Chr. okenii* (*Chr*) are indicated with arrows and algal cells (*Alg*) are denoted with triangles.

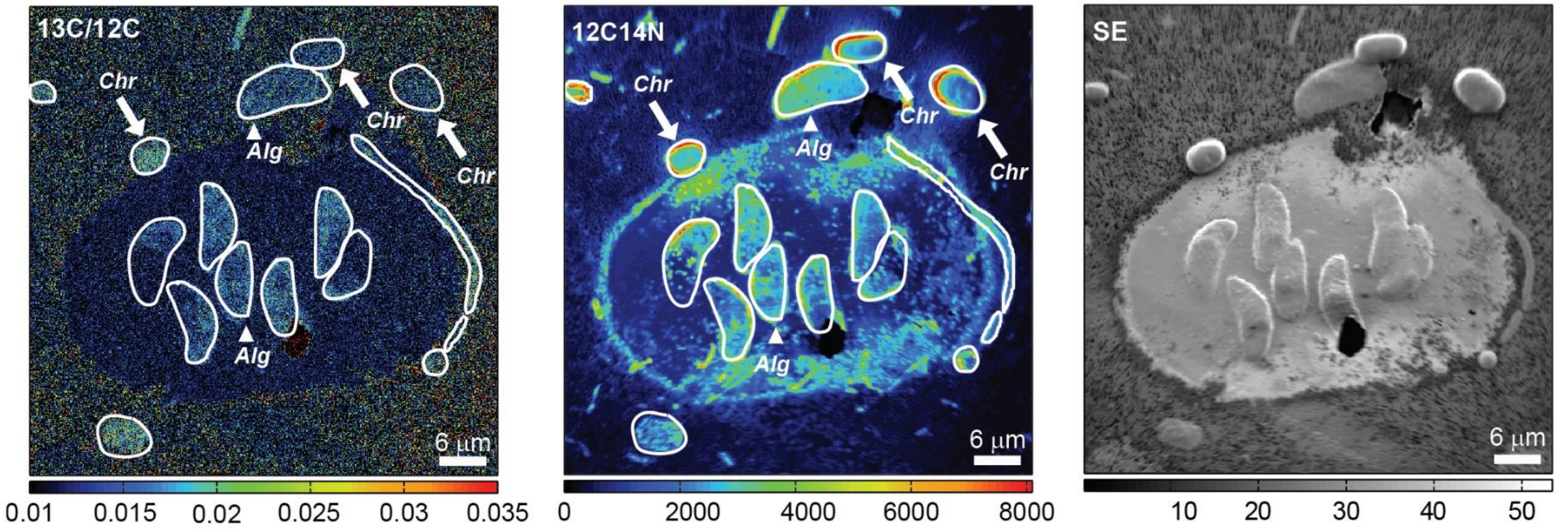

Figure S11| Motility characteristics of *Chr. okenii* in an environmental sample obtained from the chemocline during the night. (A) Trajectories of swimming *Chr. okenii* obtained from the image analyzed video micrographs, shown here for time points corresponding to 0, 4, 8 and 12 seconds of image acquisition. (B) Relative frequency distribution of swimming speeds of *Chr. okenii* under dark conditions. The mean swimming speed, 9.9 ± 2.8 μm $s^{-1}$, (mean ± SD from 180 analyzed cell trajectories) is indicated by the vertical black line.

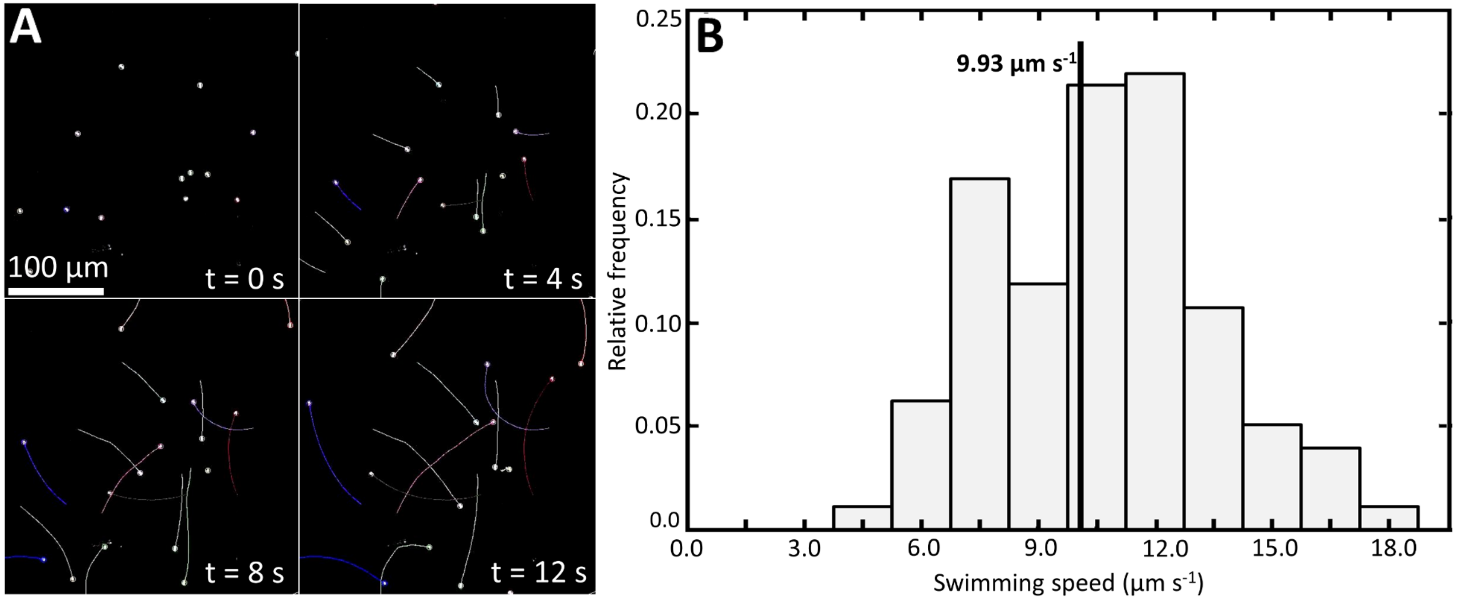

Figure S12| Contigs (>10,000 kB, min. 1 x / max. 1000 x coverage) from a combined bulk assembly of a Lake Cadagno chemocline sample and the sulfide-oxidizing enrichment culture. Size of circles represent contig size. Larger clusters and clouds signify that multiple/many contigs are located in that region. Contours of the *Chromatium okenii* MAG are highlighted in purple.

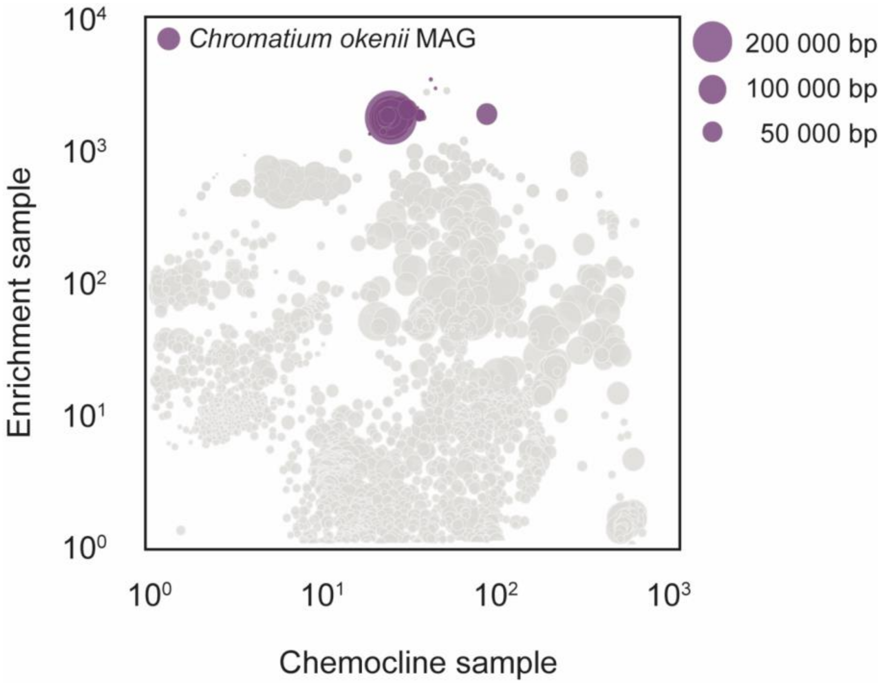

Fig S13| Phylogenetic trees showing the fraction of the diversity of (A) subunit 1 of the cytochrome cbb3-type heme-copper oxidases and (B) subunit 1 of the cytochrome bd-type recovered in the Lake Cadagno metagenome, with NorB as the outgroup. Only those sequences affiliated with bacteria known to participate in oxidative sulfur cycling are shown. *Chr. okenii* sequences from the bulk metagenome and MAG have been de-replicated and both gene IDs are shown.

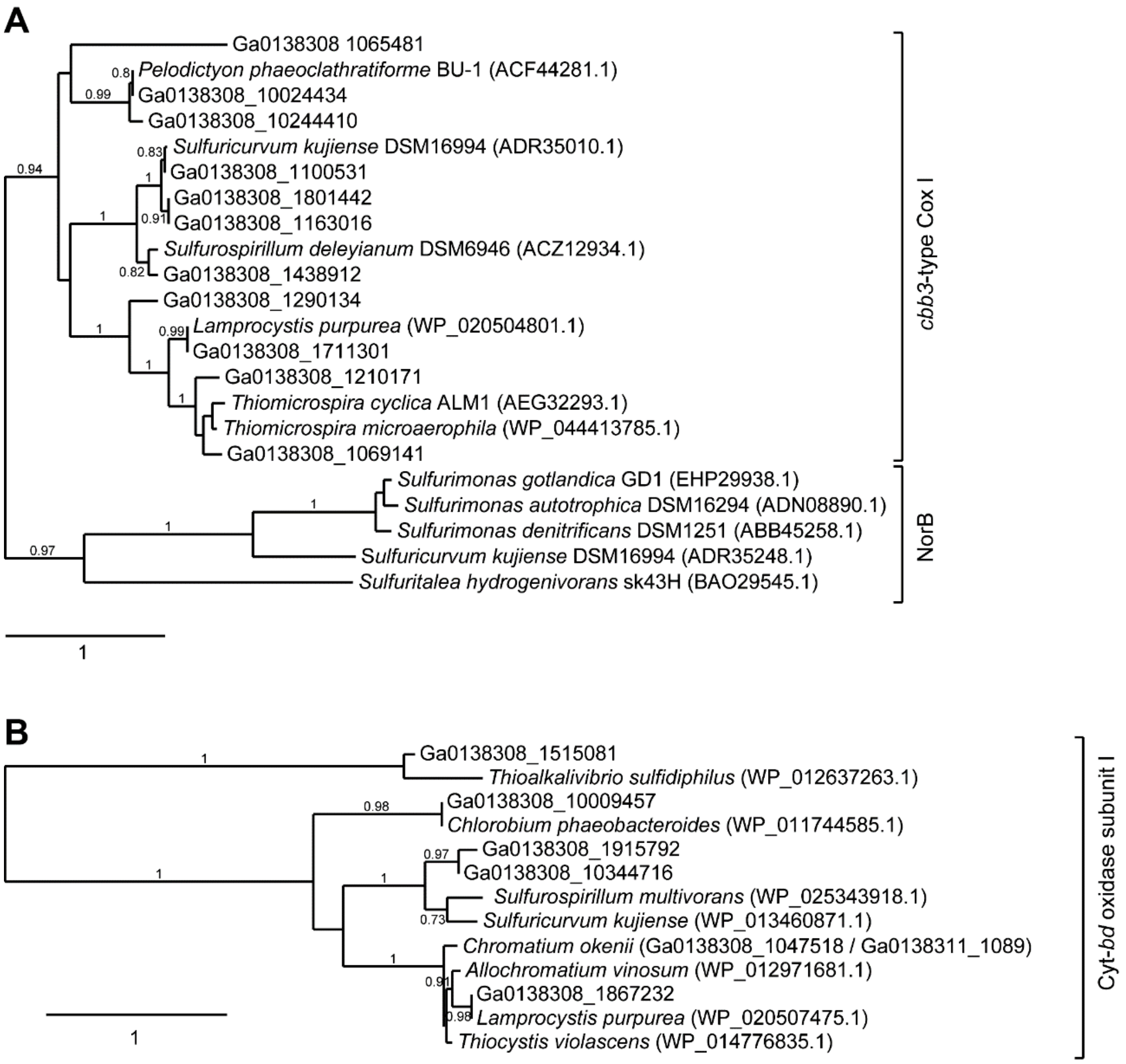

Table S1| Sequencing and assembly statistics for the metagenomics datasets recovered from the Lake Cadagno chemocline and a phototrophic, sulfide-oxidizing enrichment culture.

| **Sample** | **Chemocline** | **Enrichment Culture** |
|---|---|---|
| | *sequencing statistics* | |
| N [reads] | 184,117,826 | 84,515,200 |
| N [reads] after quality trimming | 171,779,392 (93.3%) | 81,373,588 (96.3%) |
| | *assembly statistics* | |
| Assembly size [bp] | 333,708,487 | |
| N [contigs >1000 bp] | 99,069 | |
| Max. contig size [bp] | 230,248 | |
| Mean contig size [bp] | 5,046 | |
| % GC | 49.1% | |
| Number of ORFs | 376,830 | |
